# Fat specific adipose triglyceride lipase is necessary for iron-mediated lipolysis and lipid mobilization in response to negative energy balance

**DOI:** 10.1101/2021.08.05.455308

**Authors:** Alicia R. Romero, Andre Mu, Janelle S. Ayres

## Abstract

Maintenance of energy balance is essential for the overall health of an organism. In mammals, both negative and positive energy balance are associated with disease states. To maintain their energy balance within a defined homeostatic setpoint, mammals have evolved complex regulatory mechanisms that control energy intake and expenditure. Traditionally, studies have focused on understanding the role of macronutrient physiology in energy balance. In the present study, we examined the role of the essential micronutrient iron in regulating energy balance. Using a dietary model, we found that a short course of excess dietary iron caused a negative energy balance resulting in a severe whole body wasting phenotype. This disruption in energy balance was due to an iron dependent increase in energy expenditure caused by a heightened basal metabolic rate and activity level. Using a transgenic mouse model lacking adipose triglyceride lipase (ATGL) specifically in fat tissue, we found that to meet the increased energetic demands, dietary iron caused increased lipid utilization that required fat specific ATGL-mediated lipid mobilization and wasting of subcutaneous white adipose tissue deposits. When fed dietary iron, mice lacking fat-specific ATGL activity were protected from fat wasting, and developed a severe cachectic response that is necessary to meet the increased energetic demands caused by the dietary regimen. Our work highlights the multi-faceted role of iron regulation of organismal metabolism and provides a novel *in vivo* mechanism for micronutrient control of lipolysis that is necessary for regulating mammalian energy balance.

## Introduction

Mammals have evolved complex homeostatic control mechanisms to maintain their energy balance within a defined set point. These include mechanisms that regulate nutrient acquisition, nutrient utilization and energy storage, to achieve a balance between energy intake and energy expenditure. Energy expenditure is a function of multiple factors including basal metabolism, adaptive thermogenesis and physical activity (Argilés et al., 2014; Spiegelman and Flier, 2001). Basal metabolism of an organism involves processes within the organism that are necessary to sustain life. Adaptive thermogenesis describes energy that is dissipated in the form of heat when animals consume a meal or adapts to a cold environment. Physical activity involves voluntary movements by the organism. Homeostatic control of energy balance is maintained through the interactions between genetic and environmental factors. Disruptions in these interactions that ultimately lead to energy imbalances will manifest as changes in energy storage (Spiegelman and Flier, 2001). Animals that have an energy intake that exceeds energy expenditure will have a net gain, resulting in increased energy storage. If animals have an energy intake that is less than energy expenditure, there will be a net loss, resulting in a loss of fat and other energy stores. The ability to maintain energy balance within the homeostatic set point is critical as disruptions to the balance and resulting changes in energy storage may render the organism more susceptible to diseases. For example, the increased adiposity in obesity predisposes individuals to cardiovascular disease and type 2 diabetes (O’Neill and O’Driscoll, 2015; Tune et al., 2017). Extreme causes of net energy loss can result in dangerously low body fat content that predisposes animals to infections, cardiovascular damage and musculoskeletal issues (Carrero et al., 2013; Dobner and Kaser, 2018; Rausch et al., 2021). Thus understanding the genetic and environmental factors that regulate energy balance is necessary for our understanding of the metabolic basis of disease.

The essential micronutrient iron has emerged as a critical regulator of body weight and energy balance. In children and adults, there is a greater prevalence of iron deficiency in both overweight and obese individuals (Ikeda et al., 2013). The causality of iron in the regulation of adipose tissue stores has been demonstrated in animal studies. Iron deficient diet causes increased adiposity in rats and excess dietary iron causes a reduction in body fat mass in rodents (Yook et al., 2019). The possible underlying mechanisms for iron mediated control of energy balance and the resulting changes to fat storage are likely complex and multifactorial. For example, iron status affects energy intake by regulating food consumption. Iron deficiency is associated with appetite loss that can be reversed with iron supplementation possibly through its effects on the satiety hormone leptin (Gao et al., 2015; Lawless et al., 1994; Stoltzfus et al., 2004). The critical role of iron for providing adequate levels of oxygen will control an individual’s capacity to endure the amount and rigor of physical activity. Individuals that are anemic experience fatigue, reducing the amount of physical activity they undertake. Furthermore, individuals who experience low iron without anemia have impaired adaptation to aerobic activity (Brownlie et al., 2004). Lastly, as iron regulates various aspects of energy metabolism including cellular respiration, the ability of an organism to carry out their basic basal metabolic functions will also be dependent on the iron status of the individual.

The changes in energy balance and storage controlled by the iron status of an individual will result in physiological responses involved in energy substrate mobilization, storage and breakdown. White adipose tissue (WAT) serves as the major storage depot for lipids, where excess lipid species are stored as triacylglycerol in lipid droplets. Under conditions of increased energetic demand or in response to stress stimuli, triacylglycerol is mobilized from WAT to meet organismal energy demands that are not met by diet (Duncan et al., 2007). Triacylglycerides (TG) in lipid droplets are hydrolyzed into diacylglycerol (DG) by adipose triglyceride lipase (ATGL) and monoacylglycerol (MG)/ free fatty acids and glycerol by hormone-sensitive lipase (HSL). FFA and glycerol are then released into circulation where they will be utilized by peripheral tissues. ATGL and HSL are considered essential lipolytic enzymes and are negatively regulated by insulin and IGF-1 signaling through the PKA/PKC signaling cascade (Degerman et al., 1998; Wijkander et al., 1998). *In vitro*, it was shown that iron and transferrin from serum can induce lipolysis in an acute time frame and that this is independent of PKA and PKC signaling (Rumberger et al., 2004). More recently, it was shown that the elevated levels of fatty acid mobilization, adipose tissue HSL and ATGL in obese women correlated with elevated levels of ferritin, suggesting that *in vivo*, elevated iron stores correlates with increased fatty acid mobilization (Ryan et al., 2018). While iron has an established correlative relationship with adipose tissue physiology, a causative role for iron in regulating lipid mobilization *in vivo* and how this relates to iron mediated changes in energy balance, as well as the underlying mechanisms for these processes remain unknown.

In the present study, we examined the role of the essential micronutrient iron in regulating energy balance. Using a dietary model of iron overload, we found that an acute course of excess dietary iron leads to a negative energy balance and profound whole body wasting in mice. The negative energy balance was due to iron dependent elevation in energy expenditure caused by an increase in basal metabolic rate and activity during the dark cycle, but not an increase in thermogenesis. To meet the demands of the increased energy expenditure, we found that iron caused increased lipid utilization associated with increased lipolysis and lipid mobilization, as well as fat wasting. Using a transgenic mouse model, we found that iron mediated lipolysis was dependent on ATGL activity specifically in adipose tissue, while the iron mediated increase of energy expenditure was independent of ATGL activity. Animals lacking ATGL in adipose tissue, shifted to carbohydrate utilization when fed dietary iron and exhibited a severe cachectic response due to the increased energy expenditure. Our work highlights the multi-faceted role of iron regulation of organismal metabolism and provides a novel *in vivo* mechanism for micronutrient control of lipolysis that is necessary for regulating mammalian energy balance.

## Results

### Iron rich diet causes increased energy expenditure and negative energy balance

While it is well established that iron regulates organismal metabolism and energy balance, the underlying mechanisms regulating these processes are not completely known. We established a dietary model to determine the mechanistic basis for the effects of an acute course of surplus dietary iron on organismal metabolism and energy balance. Carbonyl iron is a dietary iron supplement that is slowly absorbed and has reduced inflammatory effects on the gastrointestinal tract relative to iron sulfate-based alternatives (Devasthali et al., 2009). We fed 6-week-old C57BL/6 mice an iron rich diet (IRD) containing 2% carbonyl iron or a nutrient matched control iron diet (CID) and found that while CID-fed mice exhibited an expected slight increase in whole body weight over the course of the experiment, IRD-fed mice exhibited a drastic whole-body wasting phenotype, with mice losing approximately 20% of total body weight by day 8 post diet initiation (**Figure 1A**). This net negative energy balance caused by the IRD can be due to a decrease in energy intake and/or an increase in energy expenditure. We found that mice fed IRD eat significantly less food during the first 48 hours post diet initiation, with no significant difference in food consumption between the dietary groups by day 3 post diet initiation (**Figure 1B**). To determine if this initial food aversion response in IRD-fed mice was responsible for the whole-body wasting, we performed a pairwise feeding analysis. Pair fed mice on CID had a steep decrease in body mass within the first 48 hours of the diet regimen and increased body mass thereafter; whereas mice fed IRD *ad libitum* exhibited continual weight loss over the 8 day diet regimen (**Figure 1C**). Thus, the initial reduced feeding alone does not account for the continual whole body wasting caused by the acute course of excess dietary iron.

**Figure 1:**
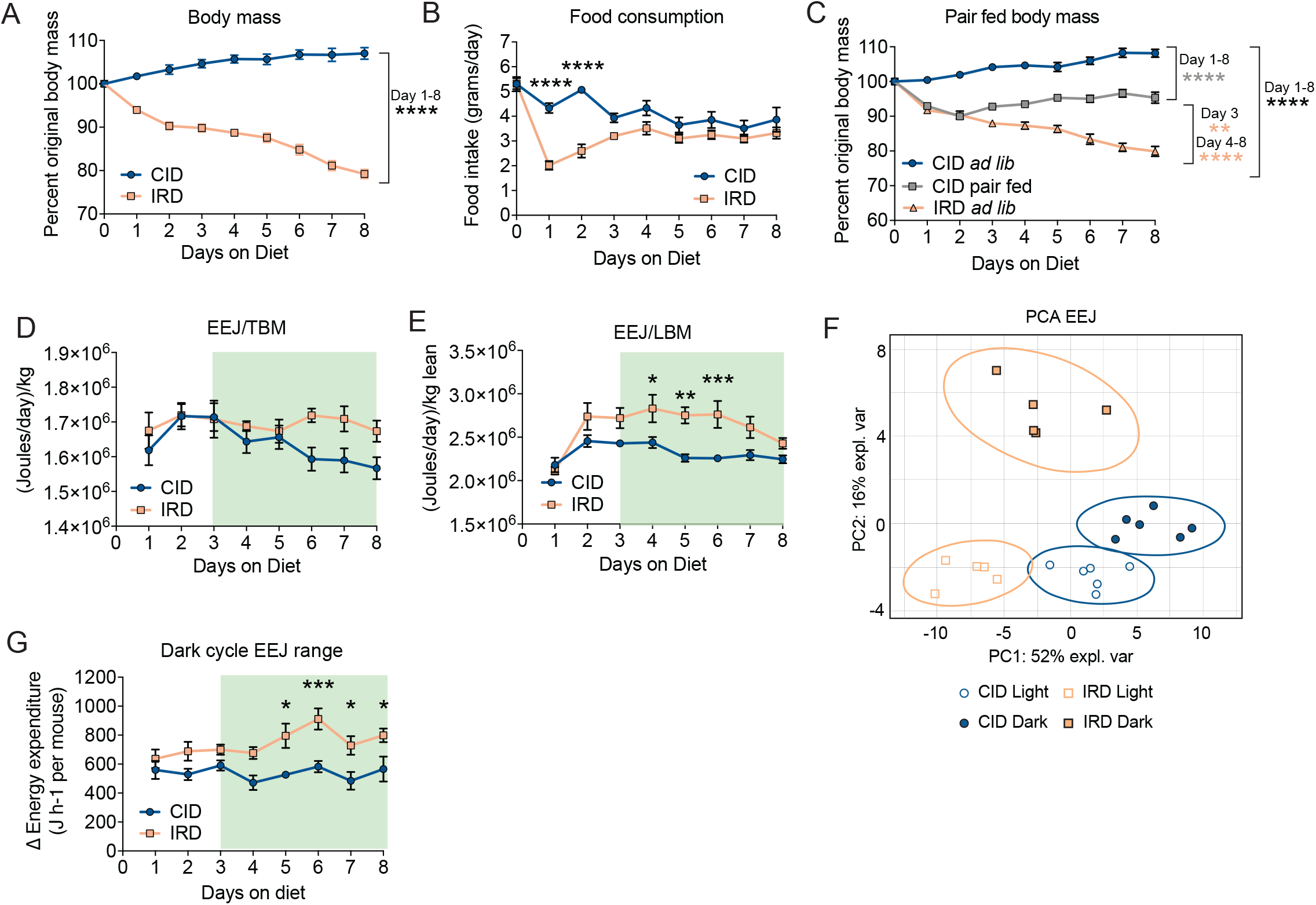
Iron rich diet causes increased energy expenditure and negative energy balance. **(A-B)** Six-week old C57BL/6 males were given control diet (CID) 2% carbonyl iron diet (IRD). Weight and food consumption were monitored. **(A)** Percent original weight over eight-day diet regimen on CID or IRD. **(B)** Average daily food consumption of mice fed CID or IRD. **(C)** C57BL/6 male mice were fed CID *ad libitum* (*ad lib)*, IRD *ad libitum* (*ad lib)*, or CID pair fed matched to average historical iron food intake values. Percent original body weight for dietary regimens shown. **(D-G)** Six-week old C57BL/6 males were housed in comprehensive laboratory animal monitoring system (CLAMS) metabolic cages for nine days. Mice were provided CID or IRD after brief acclimation period. Average daily EEJ per mouse normalized to respective daily **(D)** total body mass (TBM) and **(E)** lean body mass (LBM) on CID or IRD. Shaded region indicates period of time when animals consumed equivalent amounts of food. **(F)** Principal component analysis for CID and IRD fed mice in light/dark cycles. Ellipses are indicative of 95% confidence intervals. **(G)** The change in energy expenditure during the daily dark cycles. All CLAMS data plotted in zeitgeber time. White/black boxes on x axis represent light/dark cycles of 24-hr day. Data represent mean ±SEM; CID n=6, IRD n=5-6; *p<0.05, **p<0.01, ***p<0.001, ****p<0.0001. Related to **Supplemental Figure 1**.

To determine the effects of dietary iron on energy expenditure, we fed 6-week-old C57BL/6 mice CID or IRD for nine days and housed them in the Comprehensive Lab Animal Monitoring System (CLAMS) to measure gas exchange volumes and rates continuously over the experimental course (mice were allowed to acclimate for a day before introduction of the diets.) Mice fed an IRD displayed an increased in their energy expenditure relative to both total body mass and total lean body mass at the later stages of the dietary regimen (**Figure 1D-E** and **Supplemental Figure 1A-B**). This suggests that IRD induces an increase in total body and lean tissue energy expenditure that may account for severe iron-induced wasting. Our principal component analysis (PCA) of light/dark cycle EEJ in CID and IRD-fed mice revealed a distinct clustering of samples according to light cycle and diet. The clustering is supported by non-overlapping 95% confidence intervals(ellipses) (**Figure 1F**). IRD-fed mice were observed to exhibit a greater change in energy expenditure during their dark cycle compared to CID mice (**Figure 1G**). This increase in energy expenditure can be due to iron-induced changes in activity level, thermogenesis and/or basal metabolic rate. We observed no significant difference in total activity levels, but PCA of light/dark cycle activity levels revealed distinct clustering of dark cycle activity, suggesting that dark cycle activity level is impacted by diet (**Supplemental Figure 1C-E**). IRD fed mice exhibited a modest decrease in core body temperature, suggesting that excess dietary iron does not increase energy expenditure by increasing thermogenesis (**Supplemental Figure 1F**). Taken together, our data demonstrate that IRD induces a net negative energy balance leading to profound whole body wasting. Though reduced food intake accounts for a loss of energy intake early in the diet period, our data suggest that increased energy expenditure (mostly in the dark cycles) is responsible for driving negative energy balance thereafter and this is likely due to iron-induced changes in basal metabolic rate and dark cycle activity levels, but not adaptive thermogenesis.

### Dietary iron increases lipid utilization through lipid mobilization and wasting of fat energy stores

Negative energy balance will cause mobilization of energy substrates from endogenous stores. Mobilization of the energy stores in white adipose tissue (WAT) involves the liberation of triacylglycerol from lipid droplets stored in adipose tissue that provides free fatty acids (FFA) and glycerol to other tissues and organs. Using an *ex vivo* lipolysis assay, we found that inguinal WAT (IWAT) and gonadal WAT (GWAT) from IRD-fed mice released significantly higher levels of free non-esterified fatty acids (FFA) and free glycerol compared to mice fed CID over the course of the dietary regimen (**Figure 2A-B**). Mobilization of fat stores can result in loss of adipose tissue mass. To determine the extent of fat energy store depletion, we utilized magnetic resonance imaging (MRI) to analyze fat and lean body composition of mice fed CID or IRD for eight days. We found that the whole body wasting and increased lipid mobilization in mice fed IRD was associated with wasting of fat energy stores (**Figure 2C**). We also measured masses of representative WAT pads and found a significant reduction in the mass of IWAT, GWAT and mesenteric WAT (MWAT) in mice fed IRD (**Figure 2D-F**). MRI and direct measurement of hindlimb muscles that represent varied myofiber composition revealed that mice fed IRD also exhibit wasting of lean energy stores (**Figure 2C, G**). Further, we observed significant upregulation of two muscle-specific atrogenes, *Atrogin-1* and *Murf-1*, in gastrocnemius muscles of mice fed IRD— a hallmark of skeletal muscle wasting (**Figure 2H**) (Baehr et al., 2011; Bodine et al., 2001; Gomes et al., 2001). We also found upregulation of *Myogenin* in gastrocnemius muscles from IRD fed mice, which is associated with Type I myofiber development (**Figure 2H**) (Hughes et al., 1999). Consistent with our findings, body composition analysis on mice from our pairwise feeding experiments confirmed that fat and lean body composition were significantly reduced in the IRD *ad libitum* group relative to pair fed mice on CID (**Supplemental Figure 2A**). Similarly, we found that mice fed IRD *ad libitum* had the greatest decrease in IWAT mass and hindlimb muscle masses following after eight days on the diet regimen (**Supplemental Figure 2B-C**). In support of IRD-induced skeletal muscle wasting, we also observed the greatest upregulation of atrogene expression in gastrocnemius muscles from mice fed IRD *ad libitum* (**Supplemental Figure 2D**). Taken together, our data demonstrate that dietary iron causes lipid mobilization that results in adipose tissue wasting, as well as muscle wasting, that is independent of the initial food aversion response.

**Figure 2:**
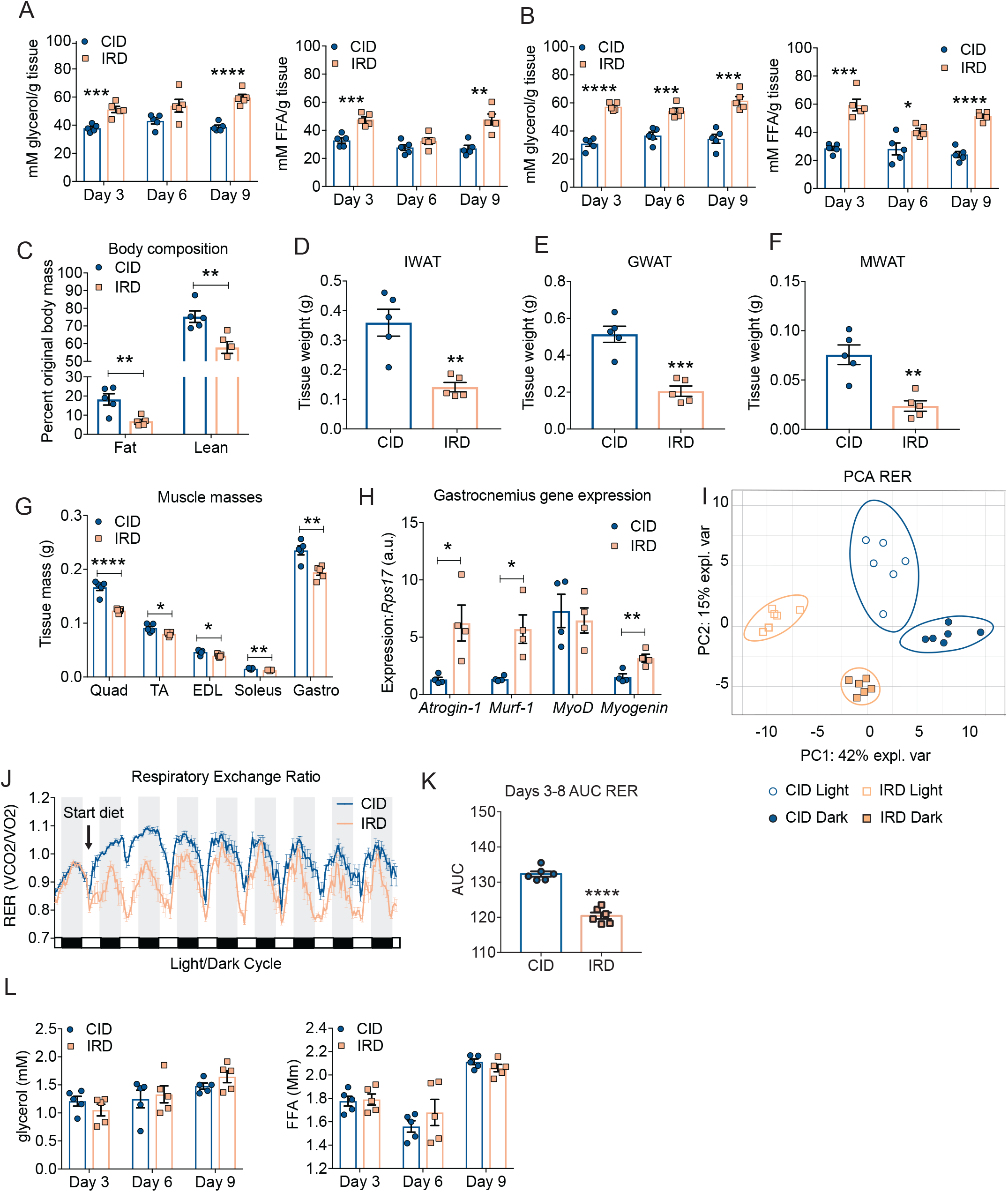
Excess dietary iron increases lipid utilization through lipid mobilization and wasting of fat energy stores. Six-week old C57BL/6 males were provided control (CID) or 2% carbonyl iron diet (IRD) for 3, 6, or 9 days. **(A-B)***Ex vivo* lipolysis assay measuring glycerol and free fatty acids (FFA) released from **(A)** IWAT **(B)** and GWAT of mice fed CID or IRD for 3, 6, or 9 days**. (C)** Body composition analyses using EchoMRI of mice fed CID or IRD for 6 days. Fat and lean mass were normalized to original body mass. **(D-F)** Tissue masses of **(D)** IWAT **(E)** GWAT and **(F)** MWAT from mice fed CID or IRD for 6 days**. (G)** Muscle masses from hindlimb of mice fed CID or IRD for 8 days (Quadricep; Quad, tibialis anterior; TA, extensor digitorum longus; EDL, soleus, and gastrocnemius; Gastro). **(H)** Gene expression in gastrocnemius of mice fed CID or IRD for 8 days **(I-K)** Mice were housed in the CLAMS and given CID or IRD. The respiratory exchange ratio over the course of the experiment was determined. **(I)** Principal component analysis for the respiratory exchange ratio (RER) CID and IRD fed mice in light/dark cycles. Ellipses are indicative of 95% confidence intervals. **(J)** Average hourly respiratory exchange ratio of mice fed CID or IRD (RER) calculated as ratio of ml CO2 OUT (respired):ml O2 IN (inhaled) **(K)** Area under the curve analysis for RER for period of matched food consumption. **(L)** Circulating levels of glycerol and FFA in serum of fasted CID or IRD mice on days 3, 6, and 9 of diet time course. Data represent mean ±SEM; CID n=4-6, IRD n=4-6; *p<0.05, **p<0.01, ***p<0.001, ****p<0.0001. Related to **Supplemental Figure 2**.

We hypothesized that the increased lipid mobilization was indicative of IRD causing a shift towards lipid utilization. To determine how IRD influences energy substrate preference, we measured respiratory exchange ratio (RER) by housing mice fed CID or IRD in the CLAMS. An RER close to 1 is indicative of carbohydrate utilization, while an RER closer to 0.7 is indicative of lipid substrate utilization (Speakman, 2013). Using PCA, we found that mice from the different dietary conditions exhibited distinct clustering during their day and night cycles, indicating that there were distinct RERs for each dietary condition and their respective light/dark cycles (**Figure 2I**). We found that IRD-fed mice exhibited a dramatic shift towards lipid utilization (**Figure 2J**). Increased lipid utilization is characteristic during periods of fasting when mice will draw from white adipose tissue (WAT) stores (Rosen and Spiegelman, 2006; Schmidt-Nielsen, 1997). We therefore performed an RER area under the curve analysis for days 3-8, when food consumption was comparable between CID and IRD-fed mice to analyze energy substrate utilization independent of differences in food consumption (**Figure 2K**). We found that the IRD induced an increase in lipid utilization independent of the fasted state. The observed increase in lipid substrate utilization was concurrent with increased relative energy expenditure and severe wasting exhibited by IRD fed mice (**Figure 1**). Despite increased release of FFA and glycerol from WAT in IRD-fed mice (**Figure 2A-B**), we observed comparable levels of circulating FFA and glycerol in mice fed either diet (**Figure 2L**), suggesting that mice fed an IRD have increased flux of circulating lipids with more rapid consumption of FFA and glycerol by peripheral tissues. Taken together, our data demonstrate that dietary iron excess causes changes to organismal energy balance, inducing a global shift towards lipid utilization that is fueled by mobilization of fat energy stores.

### An acute course of excess dietary iron does not induce insulin resistance

Iron metabolism can influence glucose homeostasis by altering insulin sensitivity. Previous work in C57BL/6 mice showed that long-term administration of dietary iron causes insulin resistance (IR) (Dongiovanni et al., 2013). We previously demonstrated in C3H/HeJ mice that an acute course of dietary iron caused transient insulin resistance during infection with an enteric pathogen (Sanchez et al., 2018). We therefore asked whether an acute course of IRD could similarly affect insulin sensitivity in C57BL/6 mice. We performed an insulin tolerance test (ITT) on days 3, 6, and 9 post-diet initiation on mice fed CID and IRD. We found no difference in insulin sensitivity between mice fed the different diets at any of the examined time points (**Figure 3A-B, Supplemental Figure 3A-D**). It has been previously demonstrated that iron overloading can lead to tissue-specific insulin resistance; notably, insulin resistance was observed in visceral WAT but not subcutaneous WAT (Dongiovanni et al., 2013; Sanchez et al., 2018). We therefore examined activation of insulin signaling in the GWAT (visceral) and IWAT (subcutaneous), as well as the liver and gastrocnemius muscle in CID and IRD fed mice. Mice fed CID and IRD for six days were injected with insulin and tissues were harvested 15 minutes post-insulin injection and processed to measure activation of AKT, a central kinase in the intracellular insulin signaling cascade (**Figure 3C**). We found no differences in phosphorylated AKT in the GWAT from mice in either dietary condition (**Figure 3D, Supplemental Figure 3E-F**). Interestingly, we found a trend towards increased phosphorylation of AKT in the IWAT, liver, and gastrocnemius muscles from mice fed IRD (**Figure 3E-G, Supplemental Figure 3G-L**). Thus, our model of acute dietary iron enrichment in C57BL/6 mice causes changes in organismal lipid physiology that is not associated with insulin resistance.

**Figure 3:**
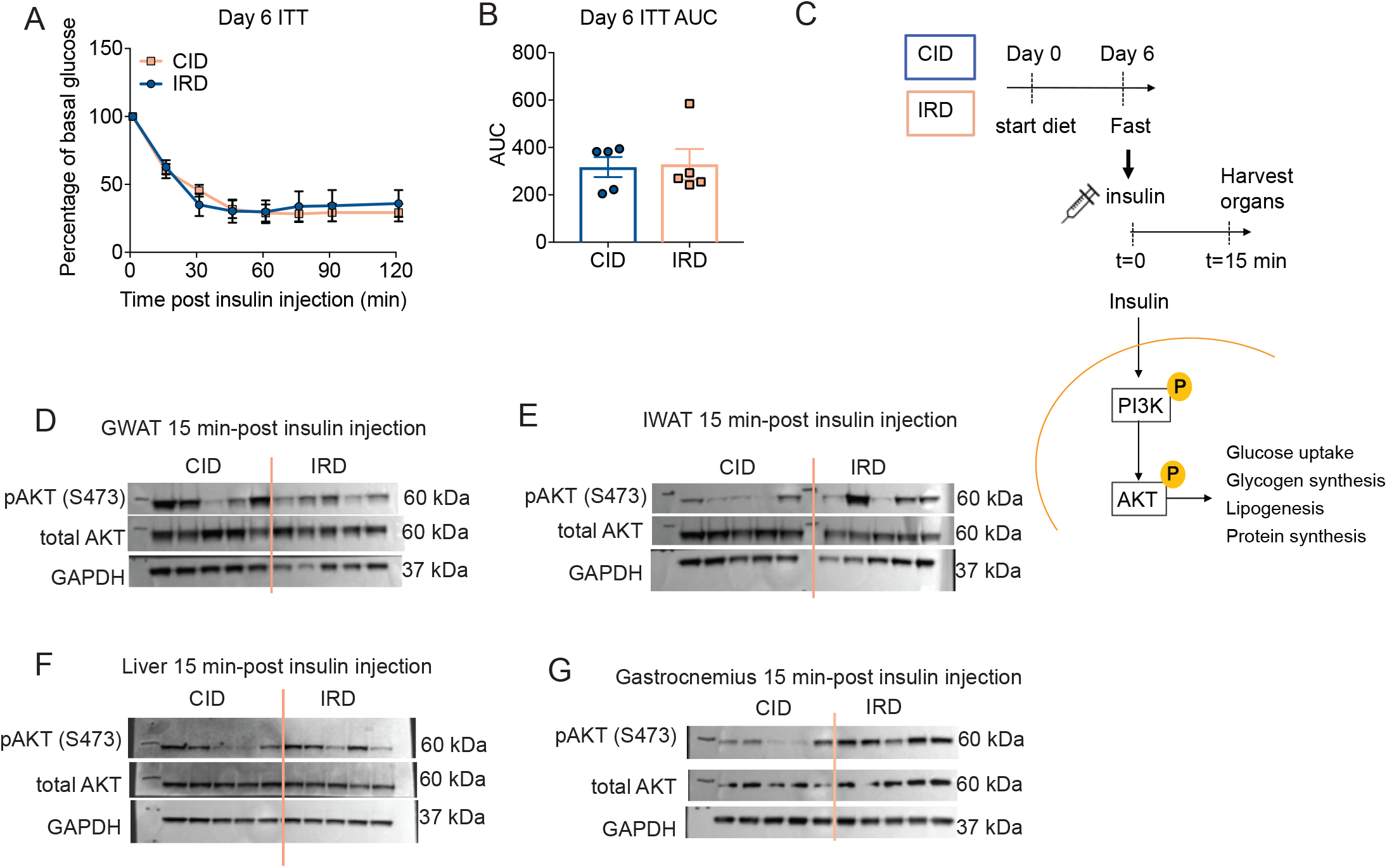
An acute course of dietary iron does not induce insulin resistance. Six-week old C57BL/6 males were provided control (CID) or 2% carbonyl iron diet (IRD) for a nine-day time course and experiments were performed on day 3, 6, and 9 of time course following a 6-hour fast. **(A-B)** Insulin tolerance test on day 6 of time course normalized to basal glucose levels taken at time of insulin injection and **(B)** area under the curve analysis of blood glucose over ITT. **(C)** Diagram for experimental setup to analyze insulin signaling. Following a 6-hour fast on day 6 of the diet time course, mice were i.p. injected with insulin and liver, fat pads, and muscle was harvested 15 minutes post insulin injection to analyze tissue-specific insulin signaling. **(D-G)** Western blot analyses for AKT protein activation in protein extracts from **(D)** GWAT, **(E)** IWAT, **(F)** liver, and **(G)** gastrocnemius muscle from hindlimb. Data represent mean ±SEM; CID n=5, IRD n=5. Related to **Supplemental Figure 3**.

### Iron-induced lipid mobilization and adipose tissue wasting is dependent on fat-specific ATGL activity

The bulk of lipid mobilization from adipose tissue is mediated through lipolysis. In canonical adipose tissue lipolysis, triglycerides stored in lipid droplets are hydrolyzed by adipose triglyceride lipase (ATGL) and hormone-sensitive lipase (HSL) to produce free glycerol and fatty acids to be supplied into circulation and fuel peripheral tissue metabolism (Haemmerle et al., 2006; Zimmermann et al., 2004). ATGL is the rate limiting lipase present on lipid droplets that is responsible for hydrolyzing triacylglycerol into diacylglycerol. We found that mice fed an IRD exhibited elevated levels of ATGL protein in both subcutaneous (IWAT) and visceral adipose tissue (GWAT) (**Figure 4A, B, Supplemental Figure 4–5)**. ATGL activity is directly regulated by its co-activator CGI-58, which is regulated by protein kinase A (PKA). Activated PKA will phosphorylate Perilipin-1 that is conjugated with CGI-58 on lipid droplet surfaces, thereby liberating CGI-58 to activate ATGL (Gruber et al., 2010; Lass et al., 2006). It was previously demonstrated that adipose tissue cultures exposed to iron rich media and human sera (transferrin rich) upregulate lipolysis in a PKA-independent manner, suggesting that iron-induced lipolysis may be regulated in a non-canonical manner, though a mechanism was not determined (Rumberger et al., 2004). Contrary to previous *in vitro* observations, we found a significant increase in the levels of phosphorylated PKAc in IWAT and GWAT of IRD fed mice, indicating that iron diet leads to increased activation of PKA in WAT *in vivo* (**Figure 4A, B, Supplemental Figure 4–5)**. Finally, IRD led to an increase in the activation of the downstream lipase, HSL, in IWAT and GWAT (**Figure 4A, B, Supplemental Figure 4–5)**. Taken together, our data demonstrate that an acute course of excess dietary iron leads to increased activation of the regulators of canonical adipose lipolysis in both subcutaneous and visceral WAT.

**Figure 4:**
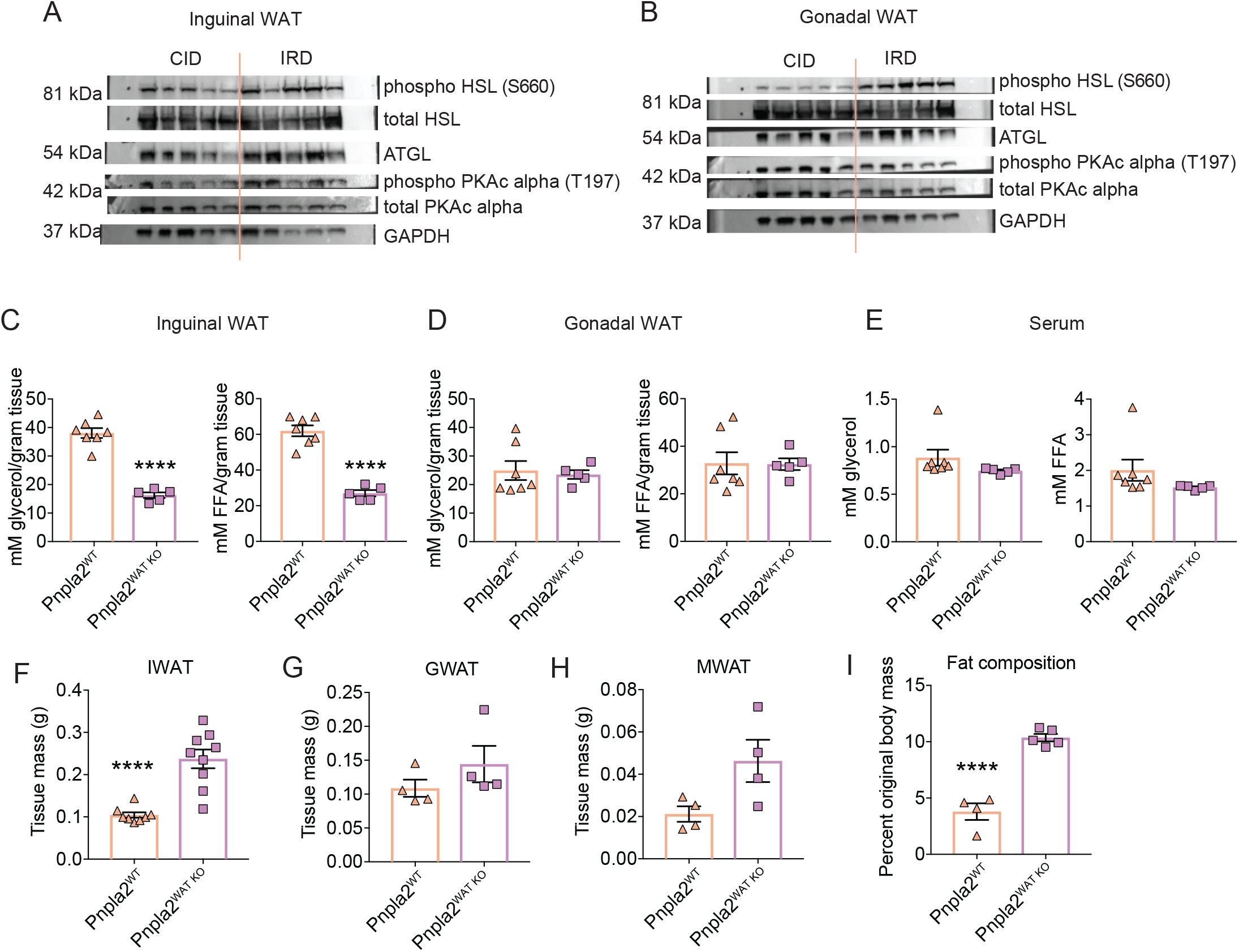
Iron induced lipid mobilization and adipose tissue wasting is dependent on fat specific ATGL activity. **(A-B)** Six-week old C57BL/6 males were given control (CID) or 2% carbonyl iron diet (IRD) for 3 days. Following a six-hour fast, mice were sacrificed and fat pads were harvested for analysis. **(A)** Western blot analysis for activated proteins in canonical lipolysis pathway in **(A)** IWAT and **(B)** GWAT protein extracts.(HSL/phospho-HSL Ser563, hormone sensitive lipase; ATGL, Adipose triglyceride lipase; PKA-c, cAMP dependent protein kinase catalytic subunit alpha; GAPDH, Glyceraldehyde3-phosphate dehydrogenase). n=5 mice per condition. **(C-D)***Pnpla2;Fabp4 cre+* (Pnpla2^WAT KO^) and *Pnpla2;Fabp4 cre-* (Pnpla2^WT^) male littermates between six and eight-weeks old were given 2% carbonyl iron diet (IRD) for 3 days and *ex vivo* lipolysis assays were performed following a 6 hour fast. Glycerol and free fatty acids (FFA) released from **(C)** IWAT **(D)** and GWAT. n=5-7 mice per condition. **(E)** Circulating levels of glycerol and FFA in serum of Pnpla2^WAT KO^ and Pnpla2^WT^ littermates fed IRD for 6 days. n=5-7 mice per condition. **(F-H)** Raw mass of fat pads from Pnpla2^WAT KO^ and Pnpla2^WT^ littermates fed IRD for five days **(F)** IWAT **(G)** GWAT **(H)** MWAT. For (F) n=8-9 mice per condition and represent two experiments combined. For (G-H) n=4 mice per condition. **(I)** Body fat composition of Pnpla2^WAT KO^ and Pnpla2^WT^ littermates fed IRD for 5 days using EchoMRI. Fat mass was normalized to original total body mass. n=4-5 mice per condition. Data represent mean ±SEM; ****p<0.0001. Related to **Supplemental Figure 4** and **5**.

To test if ATGL-dependent lipolysis in adipose tissue is necessary for mediating IRD-induced lipid mobilization and adipose tissue wasting, we generated a transgenic mouse model with the gene that encodes ATGL, *Pnpla2*, specifically deleted from adipose tissue (floxed *Pnpla2* x *Fab4 cre*). Using an *ex vivo* lipolysis assay, we found that the IWAT from IRD-fed mice lacking adipose-specific ATGL (floxed *Pnpla2;Fab4 cre+*, designated Pnpla2^WAT KO^) secreted significantly less FFA and glycerol than wild type mice fed IRD (floxed *Pnpla2;Fab4 cre-*, designated Pnpla2^WT^) (**Figure 4C**). Notably, we observed comparable levels of secreted FFA and glycerol from GWAT and no difference in circulating levels of FFA or glycerol between Pnpla2^WAT KO^ and Pnpla2^WT^ mice fed IRD (**Figure 4D,E**). The apparent difference in rates of lipid mobilization between subcutaneous and visceral adipose tissue may reflect a preferential pattern of organismal lipid storage under conditions of dietary iron excess. Consistent with this, Pnpla2^WAT KO^ mice were protected from IRD-induced IWAT wasting, but were only moderately protected from wasting of visceral fat pads, GWAT and MWAT (**Figure 4F-H)**. Though we observed differences in the pattern of subcutaneous and visceral fat pad wasting, body composition analyses showed that Pnpla2^WAT KO^ mice had significantly more total fat mass relative to Pnpla2^WT^ littermates fed IRD (**Figure 4I**). Together, these data suggest that ATGL activity is necessary for IRD-induced lipolysis, lipid mobilization and fat wasting primarily in subcutaneous adipose tissue.

### Dietary iron induced adipose-specific ATGL activity protects from wasting of lean energy stores

Having established a role for adipose tissue specific ATGL in IRD-mediated lipid mobilization and fat wasting, we next asked if ATGL activity in fat was required for the changes in energy balance and expenditure we observed in IRD-fed mice (**Figure 1–2 and Supplemental Figure 1**). Despite being protected from wasting of subcutaneous fat, Pnpla2^WAT KO^ displayed comparable total body wasting to that exhibited by their wild type littermates when fed dietary iron (**Figure 5A**). This suggests that the negative energy balance observed in IRD-fed mice occurs independent of ATGL activity in adipose tissue. In wild type mice, the negative energy balance induced by IRD is due to increased energy expenditure that is independent of differences in energy intake and instead caused by changes in the basal metabolic rate and activity levels during the dark cycle (**Figure 1** and **Supplemental Figure 1**). We found that food consumption was comparable between Pnpla2^WAT KO^ and their wildtype littermates when fed IRD (**Figure 5B**). Our analyses with the CLAMS showed that energy expenditure of Pnpla2^WT^ and Pnpla2^WAT KO^ mice was comparable under normal chow conditions (**Figure 5C**). Following introduction of IRD, Pnpla2^WT^ and Pnpla2^WAT KO^ mice showed equivalent changes in energy expenditure (**Figure 5C** and **5D**). When we examined the light/dark cycle EEJ for Pnpla2^WT^ and Pnpla2^WAT KO^ mice by PCA, we saw no influence of genotype on EEJ for light or dark cycles when animals were fed IRD (**Figure 5E**). Furthermore, we found that Pnpla2^WT^ and Pnpla2^WAT KO^ mice fed IRD displayed comparable EEJ relative to TBM over the course of the experiment (**Figure 5F**). Analysis of body temperature and activity over the course of the diet regimen revealed no difference in body temperature, total activity levels, or differences in light/dark cycle activity levels between genotypes (**Supplemental Figure 6A-D**). Taken together, our results demonstrate that ATGL activity in adipose tissue is not necessary for the effects of dietary iron on net energy balance, energy expenditure, activity levels, basal metabolism or food consumption.

**Figure 5:**
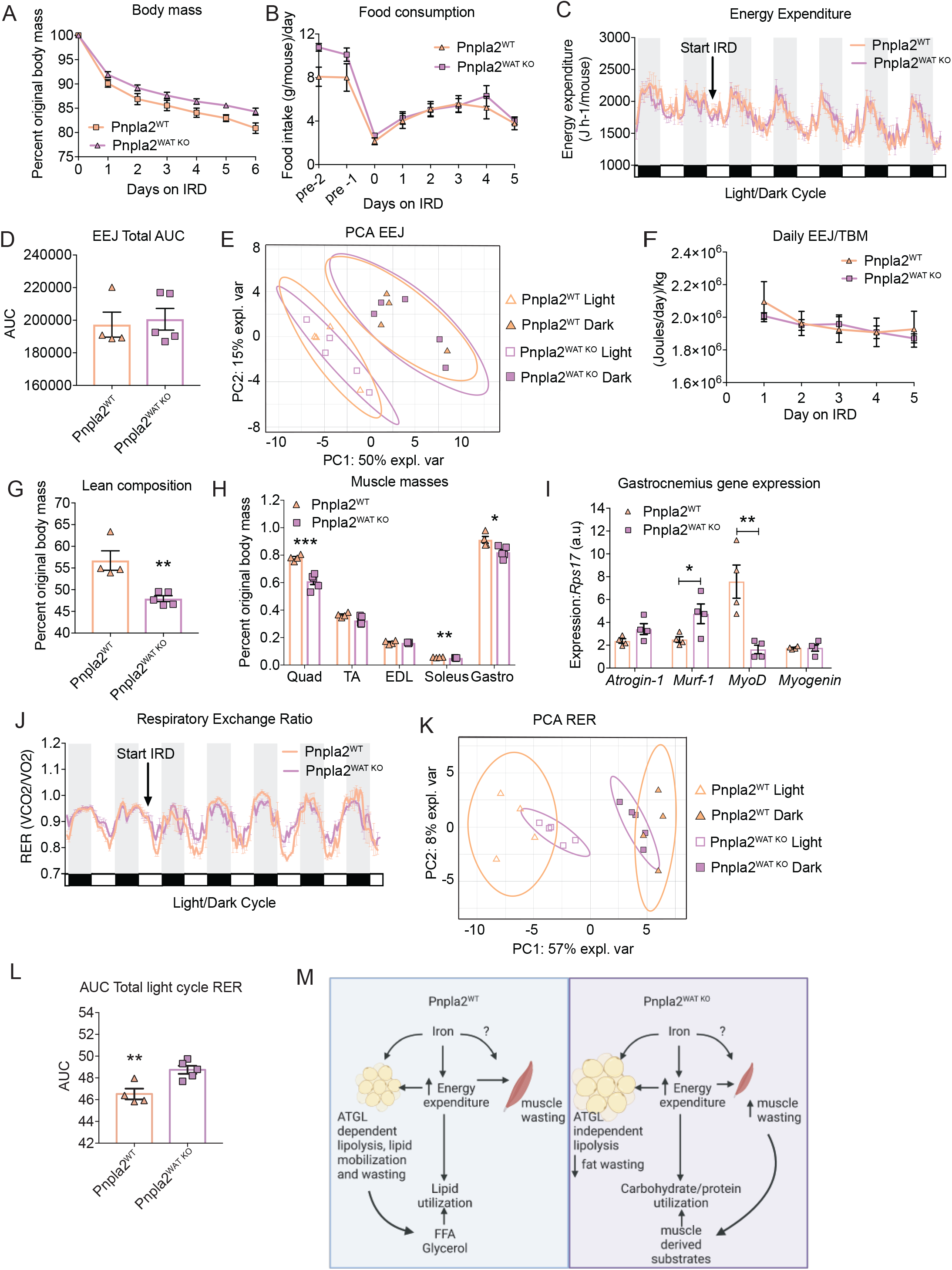
Dietary iron induced adipose-specific ATGL activity protects from wasting of lean energy stores. Littermate *Pnpla2;Fabp4 cre-* (Pnpla2^WT^) and *Pnpla2;Fabp4 cre+* (Pnpla2^WAT KO^) males between six and eight-weeks old were housed in comprehensive laboratory animal monitoring system (CLAMS) metabolic cages for 6 days. Mice were provided 2% carbonyl iron diet (IRD) after brief acclimation period. Daily measurements were taken for body mass, core temperature and food intake. **(A)** Body mass normalized to original body mass of Pnpla2^WT^ and Pnpla2^WAT KO^ mice fed IRD. n=8-9 mice per condition and represents two experiments combined. **(B)** Daily average food intake of Pnpla2^WT^ and Pnpla2^WAT KO^ mice prior to and during IRD feeding period. n=8-9 mice per condition and represents two experiments combined. **(C)** Average hourly energy expenditure (EEJ) calculated from VO2 and VCO2 (ml/hr) using the modified Weir equation and **(D)** area under the curve analysis for EEJ. n=4-5 mice per condition. **(E)** Principal component analysis of average EEJ from light/dark cycles from Pnpla2^WT^ and Pnpla2^WAT KO^ mice fed IRD. Ellipses are indicative of 95% confidence intervals. **(F)** Average daily EEJ per mouse normalized to respective daily total body mass (TBM). n=4-5 mice per condition. **(G)** Lean body composition of Pnpla2^WT^ and Pnpla2^WAT KO^ mice fed IRD using Echo MRI. Lean mass was normalized to original total body mass. n=4-5 mice per condition. **(H)** Muscle masses from hindlimb of Pnpla2^WT^ and Pnpla2^WAT KO^ mice fed IRD for five days (Quadricep; Quad, tibialis anterior; TA, extensor digitorum longus; EDL, soleus, and gastrocnemius; Gastro). n=4-5 mice per condition. **(I)** Gene expression in gastrocnemius of mice fed CID or IRD for 8 days**. (J)** Respiratory exchange ratio (RER) and **(K)** Principal component analysis of average RER from light/dark cycles from Pnpla2^WT^ and Pnpla2^WAT KO^ mice fed IRD. Ellipses are indicative of 95% confidence intervals. **(L)** Area under the curve analysis of RER for light cycle periods of CLAMS monitoring following introduction of IRD. n=4-5 mice per condition. **(M)** Model of dietary iron regelation of energy expenditure and ATGL dependent lipid mobilization. All CLAMS data plotted in zeitgeber time. White/black boxes on X axis represent light/dark cycles of 24-hr day. AUC of CLAMS analyses taken from total average values per mouse. Data represent mean ±SEM; *p<0.05, **p<0.01, ***p<0.001. Related to **Supplemental Figure 6**.

We hypothesized that Pnpla2^WAT KO^ mice undergo a compensatory increase in lean energy store wasting that can account for the whole body weight loss driven by the increased energy expenditure when these mice are fed IRD. Consistent with our hypothesis, we found that Pnpla2^WAT KO^ mice exhibited increased wasting of lean tissue stores compared to their Pnpla2^WT^ littermates fed IRD (**Figure 5G**). These differences in body composition were specific to mice fed IRD because mice lacking ATGL in adipose tissue and their wild type littermates exhibited comparable body composition when fed control chow (**Supplemental 6E**). Direct measurement of hindlimb muscle masses similarly showed that Pnpla2^WAT KO^ mice fed IRD had decreased mass of individual muscle groups compared to their wild type littermates fed IRD (**Figure 5C)**. As muscle mass is regulated by protein synthesis and degradation cascades in muscle, we analyzed expression of both atrogenes, *Atrogin-1* and *Murf-1*, and myogenesis factors, *MyoD* and *Myogenin* in gastrocnemius of IRD-fed mice. We observed increased expression of *Murf-1* and significant downregulation of *MyoD* expression in Pnpla2^WAT KO^ muscles, suggesting that both atrophic factors and lack of myogenesis signaling may contribute to enhanced muscle wasting in Pnpla2^WAT KO^ fed IRD (**Figure 5I**).

Excess dietary iron in wild type mice causes increased lipid utilization (**Figure 2**). We hypothesized that the reduced subcutaneous fat mobilization and increased skeletal muscle mobilization in IRD-fed Pnpla2^WAT KO^ mice would result in a decrease in lipid utilization. Consistent with this, we observed modest separation in RER values between Pnpla2^WAT KO^ and IRD-fed Pnpla2^WT^ mice following introduction of IRD (**Figure 5J**). Principal component analysis of the light/dark cycle RER values from IRD-fed Pnpla2^WAT KO^ and Pnpla2^WT^ mice showed that the pattern of clustering is based on the light vs. dark cycle and independent of genotype (**Figure 5K**). We therefore analyzed AUC of RER by light/dark cycles and observed a significant increase in Pnpla2^WAT KO^ RER values during the light cycle following introduction of IRD (**Figure 5L**). This suggests that during the diurnal “fasting” state, Pnpla2^WAT KO^ mice fed IRD utilized less lipid substrates relative to IRD-fed Pnpla2^WT^ littermates (**Figure 5L**). Thus ATGL activity in adipose tissue protects from lean energy store wasting by promoting lipolysis and lipid mobilization to meet the dietary iron induced increase in energy expenditure (**Figure 5M**).

## Discussion

Organismal macronutrient metabolism is intimately linked to micronutrient metabolism. While a role for iron in regulating lipid metabolism is well appreciated, the mechanistic underpinnings of these regulatory processes are unknown. Here, using an acute course of dietary iron excess, we demonstrate that iron causes profound changes in whole organismal metabolism including increased energy expenditure and lipid mobilization, resulting in a negative energy balance. We demonstrate that lipid mobilization is dependent on ATGL activity in adipose tissue and that this lipid mobilization protects from cachexia.

We found that an acute course of excess dietary iron caused an increase in energy expenditure relative to total body and lean mass, resulting in a negative energy balance. Thus our work is in agreement with clinical studies showing an association with the iron status of an individual and their energy balance as well as previous animal studies (Brownlie et al., 2004; Gao et al., 2015; Ikeda et al., 2013; Lawless et al., 1994; Stoltzfus et al., 2004; Varghese et al., 2020; Yook et al., 2019; Zhang et al., 2021). An outstanding question from our study is how the acute course of dietary iron causes an increase in energy expenditure. We propose that as iron regulates various aspects of energy metabolism, including cellular respiration, the excess dietary iron provided to animals in our study is causing increased energy expenditure by influencing their basic metabolic functions. For example, mice deficient for ferritin heavy/heart chain (FTH) gene, which encodes the ferroxidase component of the iron sequestering ferritin complex have decreased energy expenditure. This was due to hepatic and adipose tissue mitochondrial dysfunction. *Fth* is a key intracellular ferroxidase required to maintain intracellular iron homeostasis that is normally upregulated during intracellular iron overload (Blankenhaus et al., 2019). Thus, we propose one possible mechanism by which an acute course of dietary iron excess may induce increased energy expenditure is via regulation of mitochondrial homeostasis.

It has been previously reported that when fed a chronic 16-week course of dietary iron, C57BL/6 mice develop IR specifically in visceral adipose depots as well as impaired organismal insulin tolerance (Dongiovanni et al., 2013). In our study, we used an acute course of dietary iron and found that C57BL/6 mice did not develop IR in adipose tissue, skeletal muscle or liver, and also showed no changes in organismal insulin tolerance. The discrepancies between our study and that of Dongiovanni et al. may be due to the time length of the feeding regimens and suggests that in C57BL/6 mice that the IR caused by excess dietary iron may be largely influenced by length of the dietary iron regimen. In a mouse model of infectious colitis using C3H/HeJ, an acute course of dietary iron excess resulted in IR in visceral adipose tissue and changes in organismal glucose tolerance (Sanchez et al., 2018), suggesting that the ability of a short course of dietary iron to cause IR may be mouse strain specific. It is well established that inbred mouse strains have differences in iron physiology including basal differences in tissue iron content. C75BL/6 mice in particular, have a dampened iron-overloading response following introduction to an iron rich diet (Clothier et al., 1999; Dupic, 2002; Leboeuf et al., 1995; Morse et al., 1999). When considering the physiological differences in iron handling between C3H and C57BL/6 that could contribute to differential manifestations in IR development, it is important to note that C57BL/6 mice lack functional *Nramp1* while C3H have a viable *Nramp1* allele. Though *Nramp1* does not appear to be directly involved in tissue iron loading, it has been shown to be important for heme recycling and influence hepcidin expression. Failure to release heme-derived iron from macrophages dampens hepcidin signaling- leading to increased basal iron uptake. Mice lacking *Nramp1* accumulate iron in liver and spleen following erythrophagocytotic stimuli *in vivo* (Soe-Lin et al., 2009). The differences between the studies of Sanchez et al and our current work also suggest that the effects of acute excess dietary iron on IR may be dependent on the disease state of the host, and that under conditions in which homeostasis is disrupted, for example during infection, the short course of excess iron may be sufficient to induce IR. Future studies addressing the role of *Nramp1* in mediating the IR response to dietary iron in different mouse strains and an understanding of the role of acute dietary iron excess in disease states are needed.

Subcutaneous fat (SAT) is considered to have greater flux in terms of storage and mobilization capacity compared to visceral fat (VAT). Specifically, excess lipids are preferentially stored in SAT, and mobilized from SAT to accommodate negative energy balance (Luong et al., 2019; Schoettl et al., 2018). We found that an acute course of dietary iron induces wasting and mobilization of SAT and VAT indiscriminately and to a similar degree, suggesting that dietary iron excess leads to lipid mobilization events in a unconventional pattern that operates in response to negative energy balance. Our result is in line with previous reports showing that mice chronically fed an iron rich diet exhibited increased iron loading, insulin resistance and wasting of VAT compared to SAT in mice fed an iron rich diet in a chronic setting (Dongiovanni et al., 2013). Curiously, we found that mice lacking adipose-specific ATGL are protected from iron-induced fat wasting in SAT, but not VAT. We demonstrated that IWAT and GWAT have comparable levels of ATGL protein expression and activation of the PKA signaling cascade upon introduction of IRD. Therefore, the difference in fat wasting between SAT and VAT cannot be attributed to differences in ATGL expression or activation through the PKA cascade. Though PKA activation is considered the canonical signaling cascade for ATGL activation, other pathways can alter ATGL activity including PKC, MAP kinase, PI3K, NFKB and lipid peroxidation resulting from Fenton and Haber Weiss reactions (Rejholcova et al., 1988; Rumberger et al., 2004; Winterbourn, 1995; Yang and Stockwell, 2016). Indeed, in an *in vitro* study utilizing visceral adipocytes from rats, iron induced lipolysis independent of the PKA lipolytic cascade, and was proposed to occur through increased levels of lipid peroxides (Rumberger et al., 2004). In the current study, our data supports a model where iron induces lipolysis in SAT via the canonical PKA-ATGL cascade, however we propose that VAT is more sensitive to iron-derived lipolytic stimuli such as lipid peroxides or ROS.

We found that an acute course of dietary iron excess caused cachexia with IRD-fed animals exhibiting wasting of muscles with varied oxidative capacity and myofiber composition, suggesting that iron induces muscle wasting in an indiscriminate manner. Cachexia is dependent on the transcriptional upregulation of two muscle specific E3 ubiquitin ligases, *Murf-1* and *Atrongin-1* (Baehr et al., 2011; Bodine et al., 2001; Gomes et al., 2001). In agreement with this, we found that iron-induced cachexia was associated with the transcriptional upregulation of *Murf-1* and *Atrogin-1* in wasting hindlimb muscles. Iron-induced muscle wasting was more severe in IRD-fed animals lacking ATGL function in adipose tissue. We propose that this increase in the cachectic response in ATGL^WAT KO^ mice is necessary to supply muscle-derived substrates to meet the iron-induced heightened energetic demands when lipid mobilization is impaired. In support of this, we observed that ATGL^WATKO^ mice fed IRD displayed a significant shift towards carbohydrate utilization during light cycles in metabolic cages. Our finding is consistent with previous reports highlighting a diurnal pattern of WAT lipolysis driven by circadian clock transcription factors CLOCK and BMAL (both *in vivo* and *ex vivo*). Light cycles represent the diurnal murine “fasting cycle” where they preferentially utilize lipids for energy, which is dependent on circadian regulation of HSL and ATGL in WAT (Duncan et al., 2007; Shostak et al., 2013). We propose that ATGL^WATKO^ mice fed an IRD undergo increased gluconeogenesis during a prolonged period of inactivity—a process that relies on supply of muscle-derived amino acid substrates (Wu et al., 2012), to meet the increased energetic demands induced by the diet. Interestingly, from our muscle characterization we found upregulation of *Myogenin* in gastrocnemius muscles from IRD-fed mice, which is a muscle comprised of both Type I and Type II myofibers. As *Myogenin* expression is associated with Type I myofiber development, this may represent adaptive remodeling of muscles to accommodate increased reliance on lipids/beta oxidation (Hughes et al., 1999; Wu et al., 2012), that may be necessary to meet the energetic demands caused by the acute course of dietary iron excess. In summary, this work defines the mechanistic basis for dietary iron induced lipid mobilization in response to increased energy expenditure and highlights the multi-faceted role of iron regulation of organismal metabolism.

## Experimental procedures

### Mice

Six week-old male C57BL/6 mice were purchased from Jackson Laboratories and housed in our facility to acclimate for 2 days prior to experimentation. To study ATGL-dependent lipolysis, we performed crosses with *B6N.129S-Pnpla2tm1Eek/J* mice to *B6.Cg-Tg(Fabp4-cre)1Rev/J* both from Jackson labs to generate *Pnpla2;Fabp4 cre+* and *Pnpla2;Fabp4 cre-* mice. For experiments, 6-8 week old male littermates were used. All animal experiments were done in accordance with The Salk Institute Animal Care and Use Committee and performed in our AALAC-certified vivarium.

### Mouse diets and Pairwise feeding

Mice were fed a 2% carbonyl iron supplemented diet or a control diet from Envigo. Daily food intake was determined by measuring mass of food pellets daily from single-housed mice. For pair feeding experiments, iron and control diets were supplied *ad libitum* while the third Pair Fed group of mice were given control diets that were weight-matched to the historical daily food intake of a mouse given iron diet *ad libitum*.

### Body composition measurements

Total body fat and lean mass were measured using an EchoMRI machine. Total fat or total lean mass(g) was normalized to total body mass(g) to determine the percent body composition of fat and lean tissues. Fat pad masses were measured by dissecting and weighing subcutaneous fat pads (inguinal WAT), visceral fat pads (gonadal WAT), and mesenteric WAT post mortem. Muscles from hindlimb were dissected and weighed postmortem. The following muscles were harvested to represent muscles of varied myofiber composition: Quadricep (primarily fast twitch), Tibialis anterior (fast/slow twitch), extensor digitorum longus (fast/slow twitch), Soleus (slow twitch), gastrocnemius (fast/slow twitch).

### Metabolic phenotyping

Mice were singly housed in metabolic cages from the Columbus Instruments Comprehensive Lab Animal Monitoring System (CLAMS) where O_2_ consumption, CO_2_ production, and activity data were collected. Food consumption was measured by weighing food pellets daily in order to determine calorie intake. Mice were removed from metabolic cages daily to measure body mass with a scale and body composition using EchoMRI. Energy Expenditure was calculated using the modified Weir equation (modified for systems lacking urine collection):

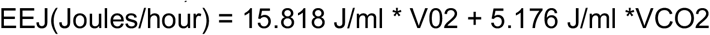

Where VO2 and VCO2 are in units of ml/hour. Respiratory Exchange Ratio (RER) was determined by dividing VCO2/VO2.

### Insulin Tolerance Tests

Animals were fasted for 6 hours in fresh cages. Blood glucose measurements were taken using a Nova Max plus by making small cuts to the tips of tails using a sterile razor blade and gently squeezing the tail from base to tip. Insulin was administered according to body mass (1U/kg) in a single intraperitoneal injection at time=0 minutes. Blood glucose measurements were taken every 15 minutes for the first hour post insulin infection and then again at 90 and 120 minutes post injection. Blood glucose measurements were normalized to their time=0 fasted blood glucose levels to generate a curve representing the percent drop in glucose levels from basal.

### Insulin Signaling Experiments and Western blot

To examine insulin signaling in tissues, mice were fasted for 6 hours and injected with insulin as described above. Injections were performed in staggered intervals. Exactly 15 minutes following insulin injection, mice were sacrificed and liver, muscle and fat pads were harvested and immediately flash frozen in liquid nitrogen. Tissues were subsequently processed for western blot analyses as described below.

### Western blot

Snap frozen tissues were ground into fine powder using ceramic mortar and pestles equilibrated in liquid nitrogen. Powder was homogenized in BeadMill24 with a ceramic bead in Tissue extraction reagent or adipose tissue protein extraction buffer with phosphatase and protease inhibitors added (Adipose tissue protein lysis buffer: 50mM Tris pH 7.5, 150mM NaCl, 1% NP-40, 0.5% Sodium Deoxycholate, 5% Glycerol). Lysates were purified by three consecutive centrifugations at 4°C to remove debris and fat and quantified using Pierce BCA protein assay kit. Samples were loaded into a 7% Tris-acetate gel following a 10-minute sonication, and a 10-minute boil at 70°C. Gels were ran for 150V for 60 minutes in Invitrogen Mini Gel Tank and transferred to nitrocellulose blot using the BioRad Turbo Transblot system for 10 minutes at 25V (1.3 A). Blots were stained with OneBlock Wes protein stain and cut into strips according to size of target protein. Strips were then washed in TBST (20mM Tris, 150mM NaCl, 0.1% Tween20 w/v) and blocked with 5% BSA in TBST for 1 hour at room temperature. Blots were incubated with primary antibodies overnight at 4°C on an orbital shaker (p-AKT, AKT, ATGL, p-HSL, HSL, p-PKAc, PKAc, GAPDH; see Catalog details in materials table). Blots were washed 3x 10 minutes with TBST and incubated with anti-rabbit for 1 hour at room temperature with Anti-Rabbit IgG HRP-linked antibodies with gentle shaking and subsequently washed 3x 10 minutes with TBST. Blots were developed and imaged using ProSignal Dura chemiluminescence in the BioRad Gel Doc XR system. Densitometry analyses were performed suing the ImageLab Software and phosphorylated proteins were normalized to total protein level and subsequently to housekeeping protein, GAPDH. Phosphorylated target proteins were probed/measured first, and the same membrane was used to measure total levels of that respective protein using the following stripping protocol: Blot was covered with stripping buffer for 5 mins and discarded. Blot was covered with fresh stripping buffer again for 5 minutes and discarded. Blot was washed two times with PBS for 10 minutes by gentle shaking. Blot was washed two times with TBST for 5 minutes by gentle shaking. After final wash, blot was blocked with 5% BSA for 1 hour at room temperature and used for subsequent antibody incubations as described above. (Mild stripping buffer (500ml): 7.5 glycine, 5 ml Tween20, 0.5g sodium dodecyl sulfate, pH 2.2)

### Gene expression analyses

Tissues were flash frozen in liquid nitrogen immediately following harvest from sacrificed animals. Tissues were ground into a powder using ceramic mortar and pestles equilibrated in liquid nitrogen. Powder was added to Trizol reagent and homogenized with a ceramic bead in a bead beater. Chloroform was added to homogenate and centrifuged to separate the organic and aqueous layers. Aqueous layer was carefully transferred to fresh nuclease-free tubes where isopropanol was added and mixture was left to precipitate at −20°C for at least 1 hour. Isopropanol/aqueous layer solution was added to a Qiagen RNeasy column and RNeasy protocol was followed from this step, including the removal of genomic DNA using Qiagen’s on-column DNase kit. RNA was eluted in nuclease-free H_2_O and quantified using a Nanodrop Spectrophotometer. cDNA was synthesized using SuperScript IV using ~200 ng RNA and oligo dT for mammalian cDNA. Real time quantitative PCR was performed using SYBR green Mix on QuantStudio 5 from Applied Biosystems. Relative standard curves method was used to analyze gene expression relative to a pooled sample dilution series. *Rps17* was used as an endogenous housekeeping control for relative normalization. Annealing temp of 60°C was used for all RT-qPCR reactions.

### *Ex vivo* lipolysis assay

Mice were fed control or 2 % carbonyl iron diets for a period of 3, 6, and 9 days and fasted in fresh cages for 6 hours. Inguinal and gonadal white adipose tissue pads were harvested, cut into 2 ~30-50 mg pieces and placed into separate wells in a 24 well plate with 300 μl of ice-cold PBS. Following the completion of tissue harvest, all fat pads were gently dabbed on a paper towel, cut with scissors in a central latitudinal location to introduce freshly cut exposed tissue, and carefully transferred to 24 well plates containing 300 μl of room-temperature Krebs-Ringer bicarbonate Hepes buffer ph 7.4 (120mM NaCl_2_, 4mM KH_2_PO_4_, 1mM MgSO_4_, 0.75 mM CaCl_2_, 30 mM Hepes pH 7.4, 10mM NaHCO_3_; solution was sterilized and stored at 4°C until day of *ex vivo* lipolysis assay when fatty acid free bovine serum albumin (to 2% wbv) and D-glucose (5mM) was added. Plates were incubated for 4 hours in a 37°C tissue culture incubator and supernatants were transferred to fresh tubes and frozen at −20°C until they were used to quantify free fatty acids and glycerol content as described below.

### Free Fatty Acid and Glycerol content assays

Serum from fasted mice and supernatant from *ex vivo* lipolysis assay were added to a 96 well plate according to the Wako Fuji Film protocol and reagents were added and incubated according to the respective FFA or glycerol Wako protocols. Standards were generated using a dilution series of glycerol or non-esterified fatty acids (WAKO). For FFA standards, we used a 1:2 dilution series from 1mM to 0.03125mM. For glycerol standards, we used the following series in mM: 8, 7, 6, 5, 4, 3, 2, 1, 0.5, 0.25. To determine FFA content, 5 μl of samples and standards were used. To determine glycerol content, 5 μl of samples were used. Following incubations with appropriate reagents, samples were quantified using a 96-well VERSAmax microplate readerand SoftMax Pro software. FFA and glycerol levels from *ex vivo* lipolysis assays were normalized to recorded tissue masses.

### Statistics

To determine the light/dark cycle and diet relationships between samples, EEJ, RER and total activity spreadsheets were transformed to represent each hour of the light/dark cycle during periods of matched food consumption. Principal component analysis was computed using the *mixOmics* R package (Rohart et al., 2017). Multivariate analysis plots were generated using the R function plotIndiv (Plot of Individuals), with the ellipses representing 95% confidence intervals of the sample space. All variable were analyzed using GraphPad Prism 7.0e. Categorical variables are expressed as numbers and percentages, and quantitative variables as means ± standard error of the mean (SEM). Where data is shown as dot plots, each dot represents a single mouse or biological replicate. One-way analysis of variance (ANOVA) with a post Tukey’s test or Two-way ANOVA were used as appropriate. Student’s two tailed t-test was performed to evaluate the differences between two independent groups or paired samples as appropriate. A *P* value <0.05 was used to reject the null hypothesis. All experiments used an *n* of 3 or more. Further information regarding specific *P* values, *n* used in each experiment, and how data are presented can be found in the figure legends.

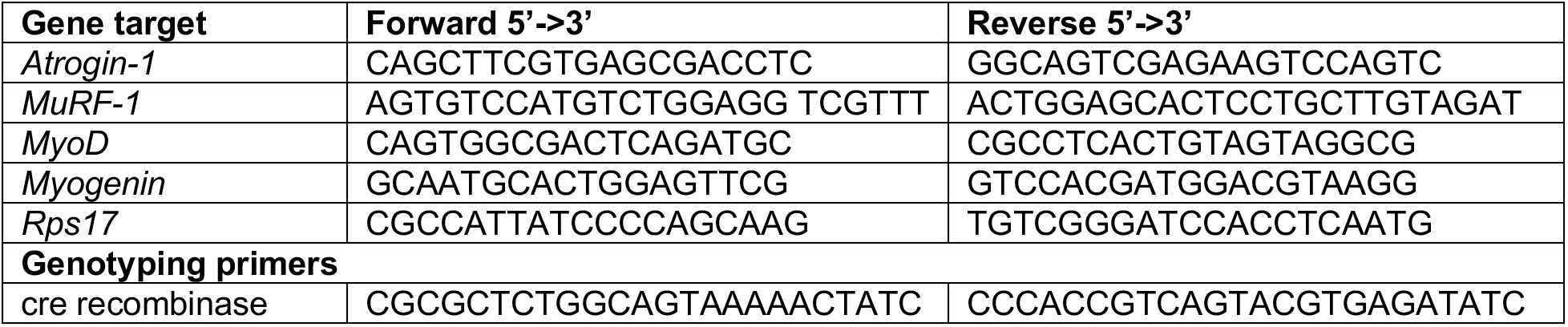

## Acknowledgements

This work was supported by an NIH awards DP1 AI144249 and R01AI114929 (JSA), and the NOMIS Foundation (JSA). Author contributions: JSA is responsible for the conceptualization, experimental design, analyzed data and wrote the paper. ARR was responsible for experimental design, analyzed the data and wrote the paper. AM performed data analysis. Figure 5M was generated using images from Biorender. JSA holds an adjunct appointment at UC San Diego.

---

| Reagent or Resource | Source | Identifier |
| --- | --- | --- |
| **Antibodies** |  |  |
| Rabbit mAb AKT | Cell Signaling | Cat#:9272S |
| Rabbit mAB Phospho-AKT (Ser473)(193H12) | Cell Signaling | Cat#:4058S |
| Rabbit mAb GAPDH XP (D16H11) | Cell Signaling | Cat#:2118S |
| Anti-Rabbit IgG HRP-linked Ab | Cell Signaling | Cat#: 7074S |
| Rabbit mAB ATGL | Cell Signaling | Cat#:2439S |
| Rabbit mAB phospho-HSL(Ser660) | Cell Signaling | Cat#:4126S |
| Rabbit mAB HSL | Cell Signaling | Cat#: 18381T |
| Rabbit mAB PKA-C | Cell Signaling | Cat#: 5842T |
| Rabbit mAB phospho-PKA-C (Thr197) | Cell Signaling | Cat#: 5661T |
| **Chemicals, reagents, peptides and recombinant proteins** |  |  |
| Tissue extraction reagent II | Invitrogen | Cat#:FN0081 |
| Trans-Blot® Turbo™ Midi 0.2 µm Nitrocellulose Transfer Packs | BioRad | Cat# 1704159 |
| SYBR Green | BioRad | Cat# 1725125 |
| Humulin R regular insulin u-100 HI-210 | Lily | Cat# 07-843-8053 |
| Halt Protease and Phosphatase inhibitor cocktail (100x) | Fischer Scientific | Cat# 78440 |
| Nupage Tris-Acetate SDS Running Buffer | ThermoFischer Scientific | Cat# LA0041 |
| Nupage 7% Tris-Acetate Gel 1.0mm X 12 well | ThermoFischer Scientific | Cat#EA03552BOX |
| Nupage LDS Sample Buffer (4x) | Invitrogen | Cat#: NP0007 |
| Protease Inhibitor Cocktail (100x) | Sigma | Cat# P8340 |
| Ponceau | Sigma | Cat#:G7021 |
| D-(+)-Glucose | Sigma | Cat#:G7021 |
| Sodium Chloride | Fisher Chemical | Cat#: S271-3 |
| Dulbecco's Phosphate Buffered Saline | Gibco | Cat#: 14190-144 |
| Bovine Serum Albumin, heat shock treated | Fisher Bioreagents | Cat#:BP1600100 |
| Tris, Hydrochloride, ULTROL Grade | Millipore Sigma | Cat#:1185-53-1 |
| Tween20 | Sigma-Aldrich | Cat#:P1379-500ml |
| Potassium Phosphate Monobasic | Fischer Scientific | Cat#:P285-500 |
| Magnesium Sulfate | Fischer Scientific | Cat#: M63-500 |
| Calcium Chloride | Sigma | Cat#:C1016-500 |
| Hepes | Sigma | Cat#:3375 |
| Sodium Bicarbonate | Tocris Bioscience | Cat#:3152 |
| Albumin, Bovine Serum, Low Fatty Acid, Low Endotoxin, Low IgG (NZ) (BSA) | US Biological Life Sciences | Cat#: A1311 |
| Sodium Dodecyl Sulfate (SDS), Lauryl | ThermoFischer Scientific | Cat#: 28365 |
| Glycine | Millipore Sigma | Cat#: 410225-250G |
| Glycerol | Fischer Scientific | Cat#: BP229-1 |
| UltraPure distilled water | Invitrogen | Cat#: 10977-015 |
| Sodium Deoxycholate | Sigma | Cat#: D6750-100 |
| NP-40 | Sigma | Cat#: 492018-50ML |
| **Critical Commercial Assays** |  |  |
| Superscript IV Reverse Transcriptase | Invitrogen | Cat# 18090010 |
| RNase OUT | Invitrogen | Cat# 10777019 |
| Trizol Reagent | Invitrogen | Cat# 15596026 |
| RNeasy Mini Kit | Qiagen | Cat# 74104 |
| RNase-Free DNase Set | Qiagen | Cat# 79254 |
| Wako HR series NEFA-HR(2) Color reagent A | WAKO Fuji Film | Cat# 999-34691 |
| Wako HR Series NEFA-HR(2) Solvent A | WAKO Fuji Film | Cat# 995-34791 |
| Wako HR Series NEFA-HR(2) Color Reagent B | WAKO Fuji Film | Cat# 991-34891 |
| Wako HR Series NEFA-HR(2) Solvent B | WAKO Fuji Film | Cat# 993-35191 |
| Free Glycerol Reagent | Sigma | Cat# F6428-40ml |
| OneBlock Wes protein stain | Prometheus Protein Biology Products | Cat# 20-311 |
| Pierce BCA Protein Assay Kit | Fischer Scientific | Cat# 23225 |
| ProSignal Dura | Prometheus Protein Biology Products | Cat# 20-301 |
| **Experimental models: organisms/strains** |  |  |
| Mouse: C57BL/6 | The Jackson Laboratory | 000664 |
| Mouse: B6.Cg-Tg(Fabp4-cre)1Rev/J | The Jackson Laboratory | 005069 |
| Mouse: B6N.129S-Pnpla2tm1Eek/J | The Jackson Laboratory | 02478 |
| **Diets** |  |  |
| 2% carboyl iron enriched chow | Envigo | Cat# TD.08714 |
| Control iron chow | Envigo | Cat# TD.99398 |
| **Equipment and software** |  |  |
| Ohaus® - CS200 - Portable Scale (discontinued) | Ohaus | CS200 |
| VERSAmax Microplate reader | Molecular devices |  |
| SoftMax Pro 5.4.5 Microplate Data Acquisition and Analysis Software | Molecular devices |  |
| Nova Max glucose test strips | Novacares | Cat# 43437 |
| Nova Max plus glucose meter | Novacares | Cat# 43435 |
| EchoMRI™-100H Body Composition Analysis | EchoMRI |  |
| BeadMill24 benchtop bead-based tissue homogenizer | Fischer Scientific | 15-340-163 |
| QuantStudio5 Real-Time 384-well | Applied Biosystems |  |
| QuantStudio5 Design and Analysis software v1.5.0 | Applied Biosystems |  |
| Gel doc XR +Gel documentation System | BioRad | 1708195 |
| Image Lab Software 5.2.1 | BioRad |  |
| Digi-sense Type J/K/T Thermocouple meter | Digi-sense | Cat#:20250-91 |
| NanoDrop™ One/OneC Microvolume UV-Vis Spectrophotometer | ThermoFisher Scientific | ND-ONE-W |
| CLAMS | Columbus Instruments |  |
| BD Microtainers SST | BD | Cat#:202204030 |
| Mini Gel Tank | Invitrogen | Cat#: A25977 |

## Supplemental Figure Legends

**Supplemental Figure 1:**
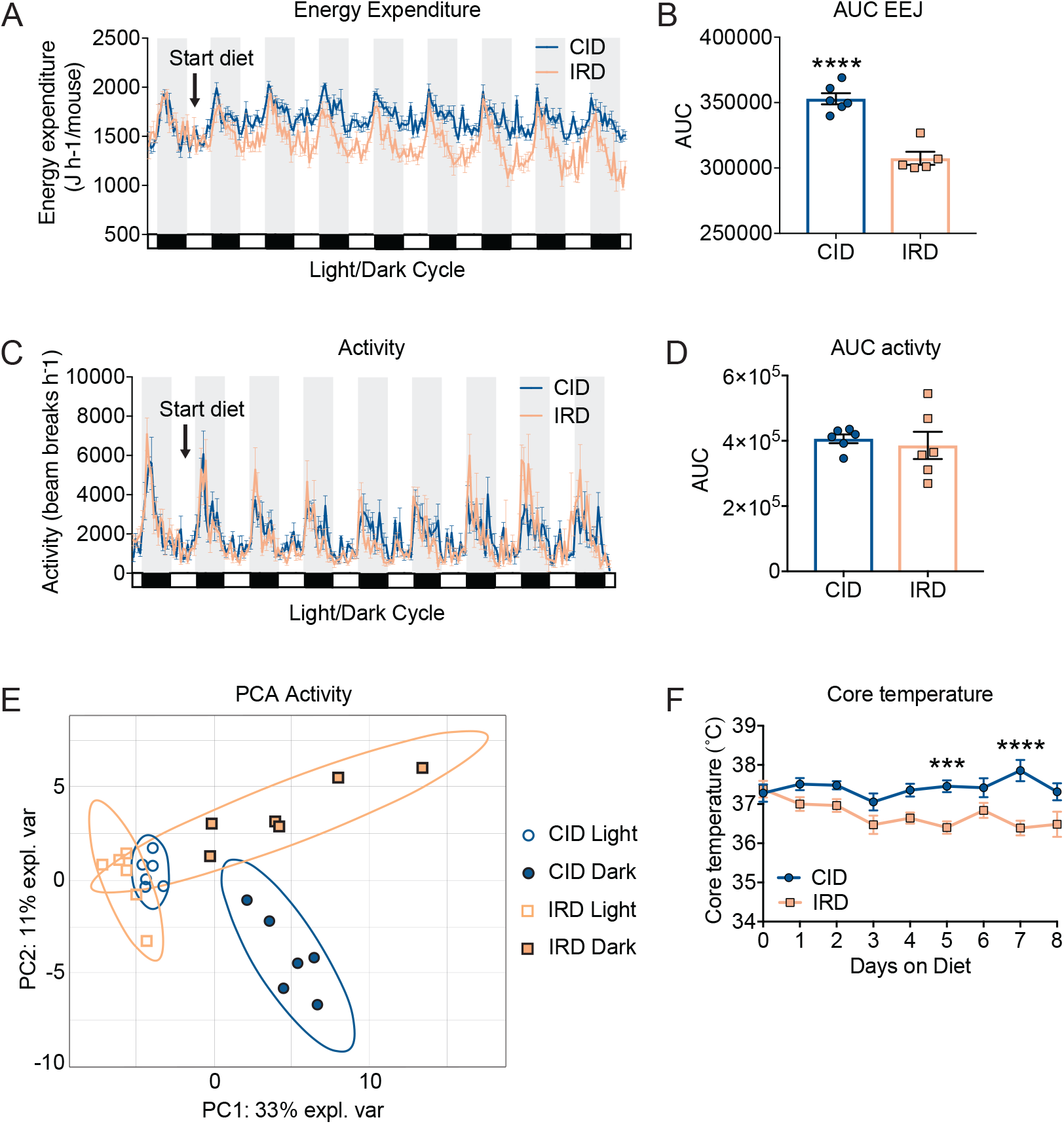
Iron rich diet causes increased energy expenditure and negative energy balance. Six-week old C57BL/6 males were housed in comprehensive laboratory animal monitoring system (CLAMS) metabolic cages for 9 days. Mice were provided control (CID) or 2% carbonyl iron (IRD) diet after brief acclimation period. Daily measurements were taken for body mass, core temperature and food intake at the same time of day. **(A)** Average hourly energy expenditure for mice fed CID or IRD (EEJ) calculated from VO2 and VCO2 (ml/hr) using the modified Weir equation. **(B)** Area under the curve analysis for EEJ data in 1D over total time in CLAMS. **(C)** Average hourly activity levels plotted as total X directional beam breaks per hour for mice fed CID or IRD in CLAMS. **(D)** Area under the curve analysis for total activity in CLAMS. **(E)** Principal component analysis for activity of CID and IRD fed mice in light/dark cycles. Ellipses are indicative of 95% confidence intervals. **(F)** Daily core temperature of CID or IRD fed mice measured with rectal thermometer. All CLAMS data plotted in zeitgeber time. White/black boxes on X axis represent light/dark cycles of 24-hr day. Data represent mean ±SEM; For **(A-E)** CID n=6, IRD n=6; For **(F)** n=11 for both diets and are two experiments combined; ***p=0.0009, ****p<0.0001. Related to **Figure 1**.

**Supplemental Figure 2:**
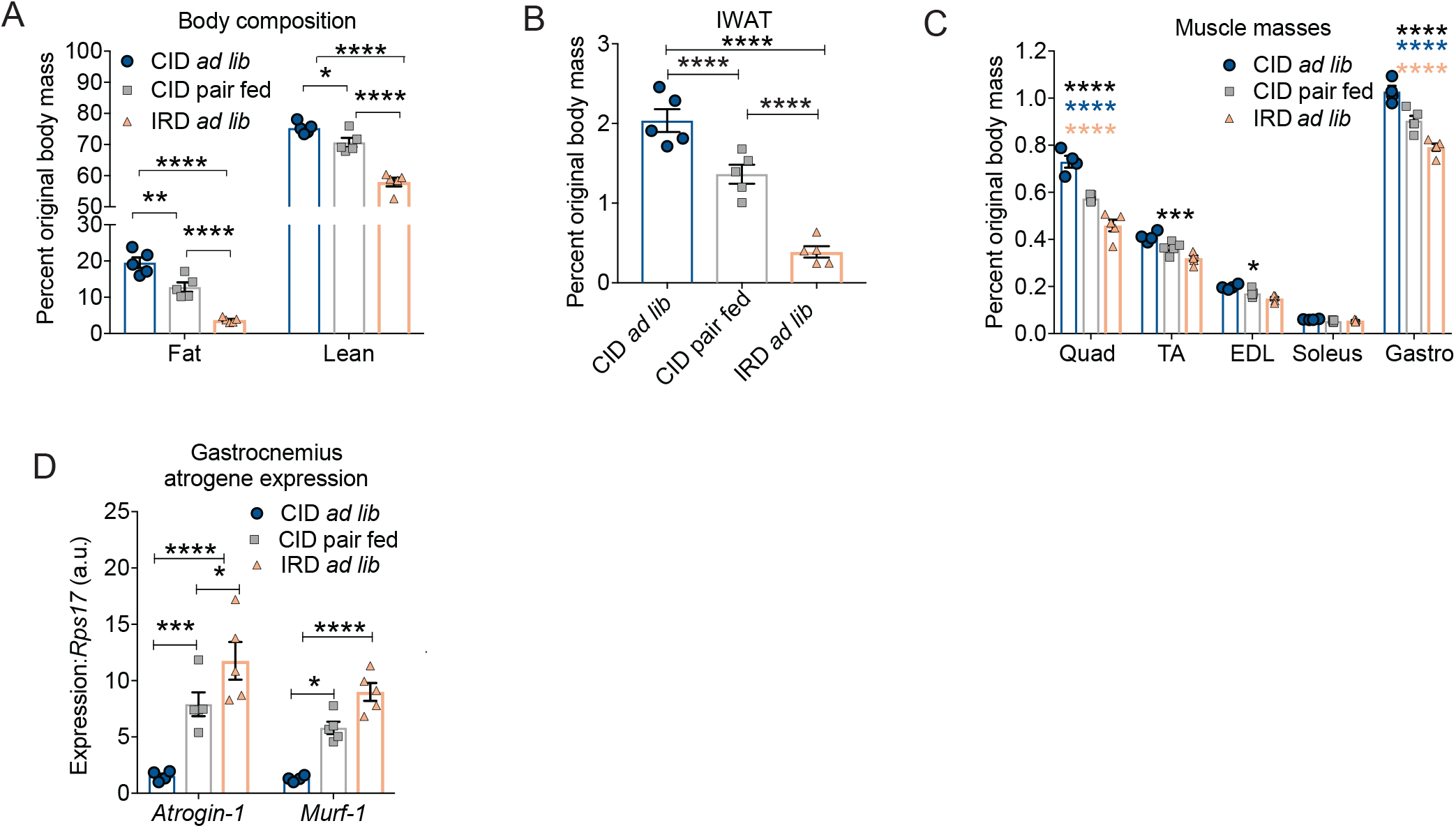
Excess dietary iron increases lipid utilization through lipid mobilization and wasting of fat energy stores. Six-week old C57BL/6 males were provided control (CID) or 2% carbonyl iron diet (IRD). Daily measurements were taken to determine food intake. Pairwise feeding experiment were done. CID *ad libitum* (CID *ad lib*), IRD *ad libitum* (IRD *ad lib*) and CID matched to the average daily food intake of iron diet mice according to historical daily food intake (CID pair fed). **(A)** Body composition analyses on day 8 of diet regimen using Echo MRI. Fat and lean mass were normalized to original body mass. **(B)** IWAT fat pad mass on day 8 of diet regimen normalized to original body mass. **(C)** Muscle masses on day 8 of diet regimen normalized to original body mass. Blue asterisks indicate comparisons between CID *ad lib* and CID pair fed; Peach asterisks indicate comparisons between IRD *ad lib* and CID pair fed; Black asterisk indicate comparisons between CID *ad lib* and IRD *ad lib*. (Quadricep; Quad, tibialis anterior; TA, extensor digitorum longus; EDL, soleus, and gastrocnemius; Gastro). **(D)** Atrogene expression in gastrocnemius muscles. Data represent mean ±SEM; CID *ad lib* n=5, IRD *ad lib* n=5, CID pair fed n=5; *p<0.05, **p<0.01, ***p<0.001, ****p<0.0001. Related to **Figure 2**.

**Supplemental Figure 3:**
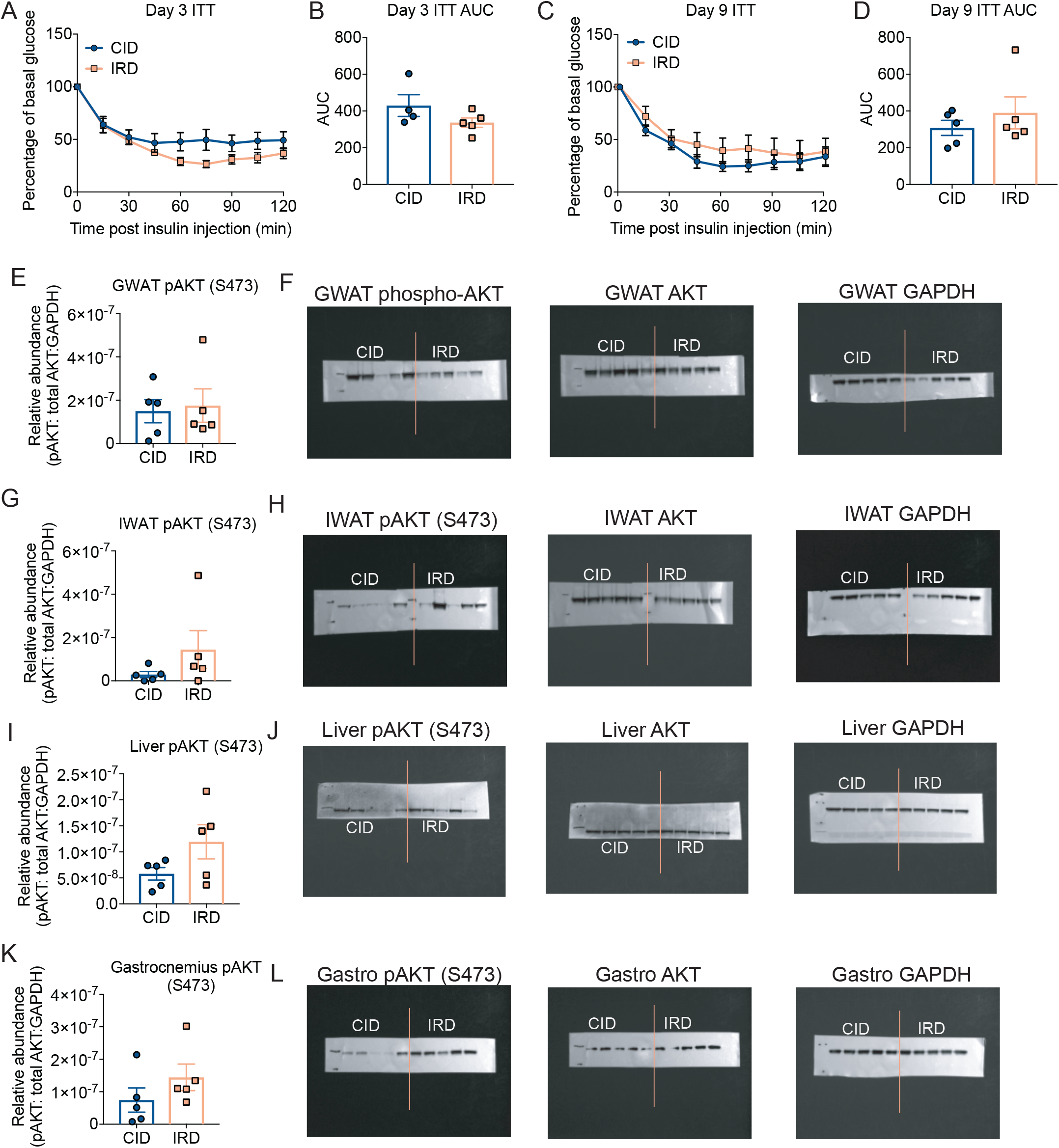
An acute course of dietary iron does not induce insulin resistance. Six-week old C57BL/6 males were provided control (CID) or 2% carbonyl iron diet (IRD) for a nine-day time course and experiments were performed on day 3, 6, and 9 of time course following a 6-hour fast. **(A-B)** Insulin tolerance test (ITT) on day 3 **(A)** ITT curve; **(B)** area under the curve analysis of **(A)**; **(C-D)** Insulin tolerance test on day 9 **(C)** ITT curve; **(D)** area under the curve analysis of **(C)**. For **(A)** and **(C)** normalized to basal glucose levels taken at time of insulin injection. **(E)** Densitometry analyses for AKT activation for GWAT blots from **Figure 3D**. **(F)** Original blots for **Figure 3D**. **(G)** Densitometry analyses for AKT activation for IWAT blots from **Figure 1E**. **(H)** Original blots for **Figure 3E**. **(I)** Densitometry analyses for AKT activation for liver blots from **Figure 3E**. **(J)** Original blots for **Figure 3F**. **(K)** Densitometry analyses for AKT activation for gastrocnemius blots from **Figure 3G**. **(L)** Original blots for **Figure 3G**. Data represent mean ±SEM; CID n=5, IRD n=5. Related to **Figure 3**.

**Supplemental Figure 4:**
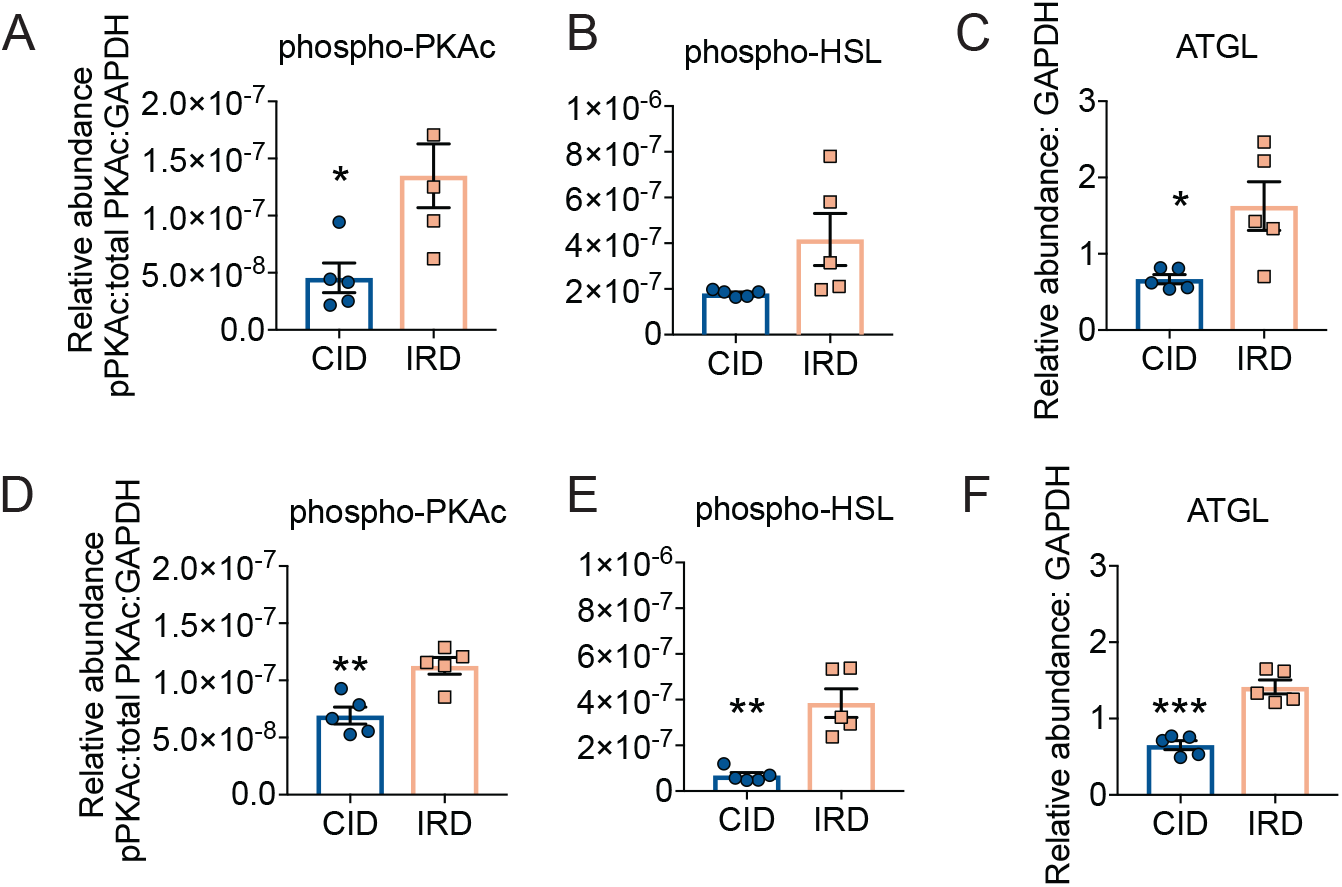
Iron induced lipid mobilization and adipose tissue wasting is dependent on fat specific ATGL activity. **(A-C)** Densitometry quantification of activated proteins in lipolysis pathway from IWAT (**Figure 4A**) normalized to respective total protein and/or housekeeping GAPDH protein. **(D-F)** Densitometry quantification of activated proteins in lipolysis pathway from GWAT (**Figure 4B**) normalized to respective total protein and/or housekeeping GAPDH protein. Data represent mean ±SEM; CID n=5, IRD n=5; *p<0.05, **p<0.01, ***p<0.001. Related to **Figure 4**.

**Supplemental Figure 5:**
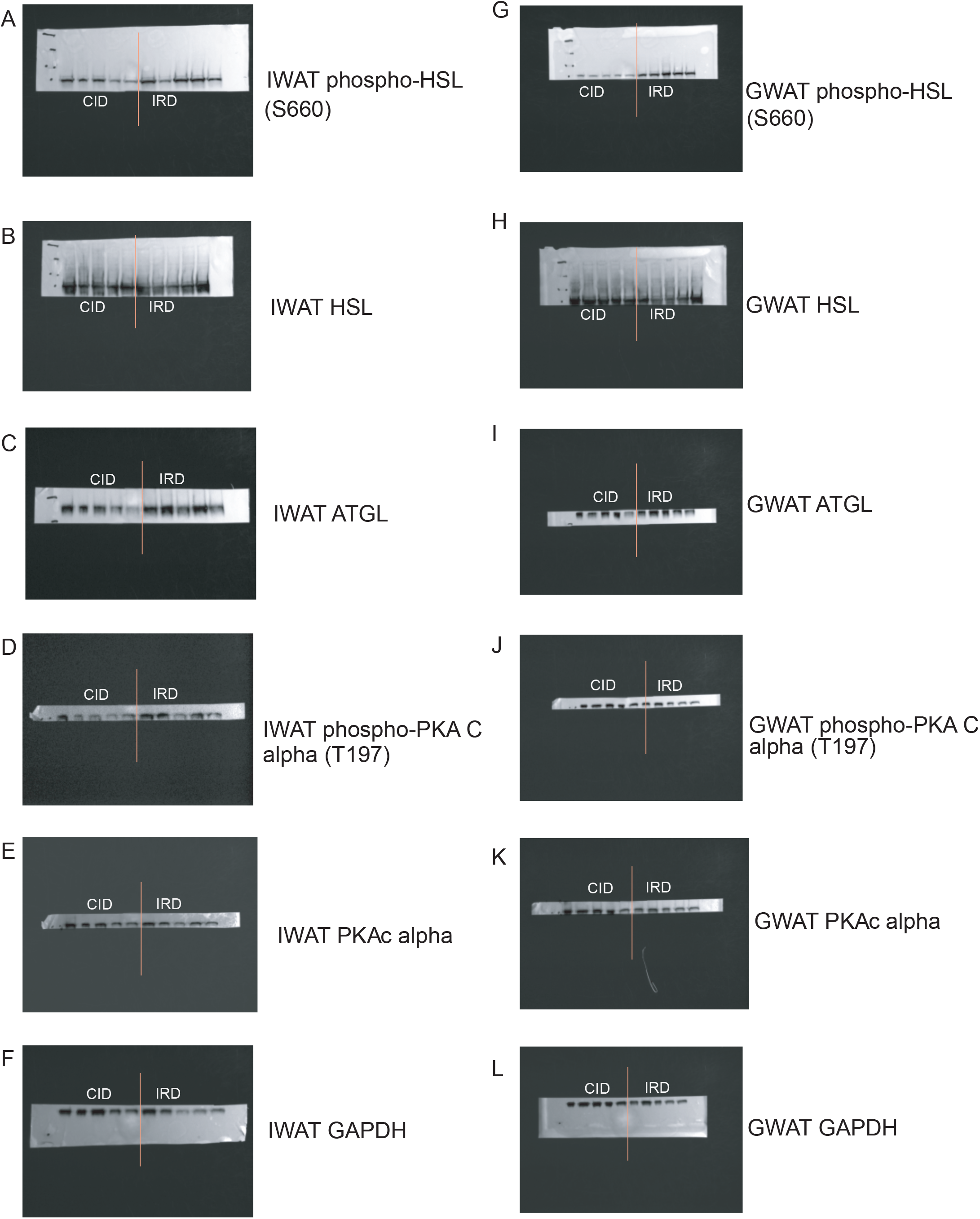
Iron induced lipid mobilization and adipose tissue wasting is dependent on fat specific ATGL activity. **(A-F)** Uncropped Western blot imaging for **Figure 4A** protein extracts from IWAT. (HSL/phospho-HSL Ser563, hormone sensitive lipase; ATGL, Adipose triglyceride lipase; PKA-c, cAMP dependent protein kinase catalytic subunit alpha; GAPDH, Glyceraldehyde3-phosphate dehydrogenase). **(G-L)** Uncropped Western blot imaging for **Figure 4B** protein extracts from GWAT. (HSL/phospho-HSL Ser563, hormone sensitive lipase; ATGL, Adipose triglyceride lipase; PKA-c, cAMP dependent protein kinase catalytic subunit alpha; GAPDH, Glyceraldehyde3-phosphate dehydrogenase). Related to **Figure 4**.

**Supplemental Figure 6:**
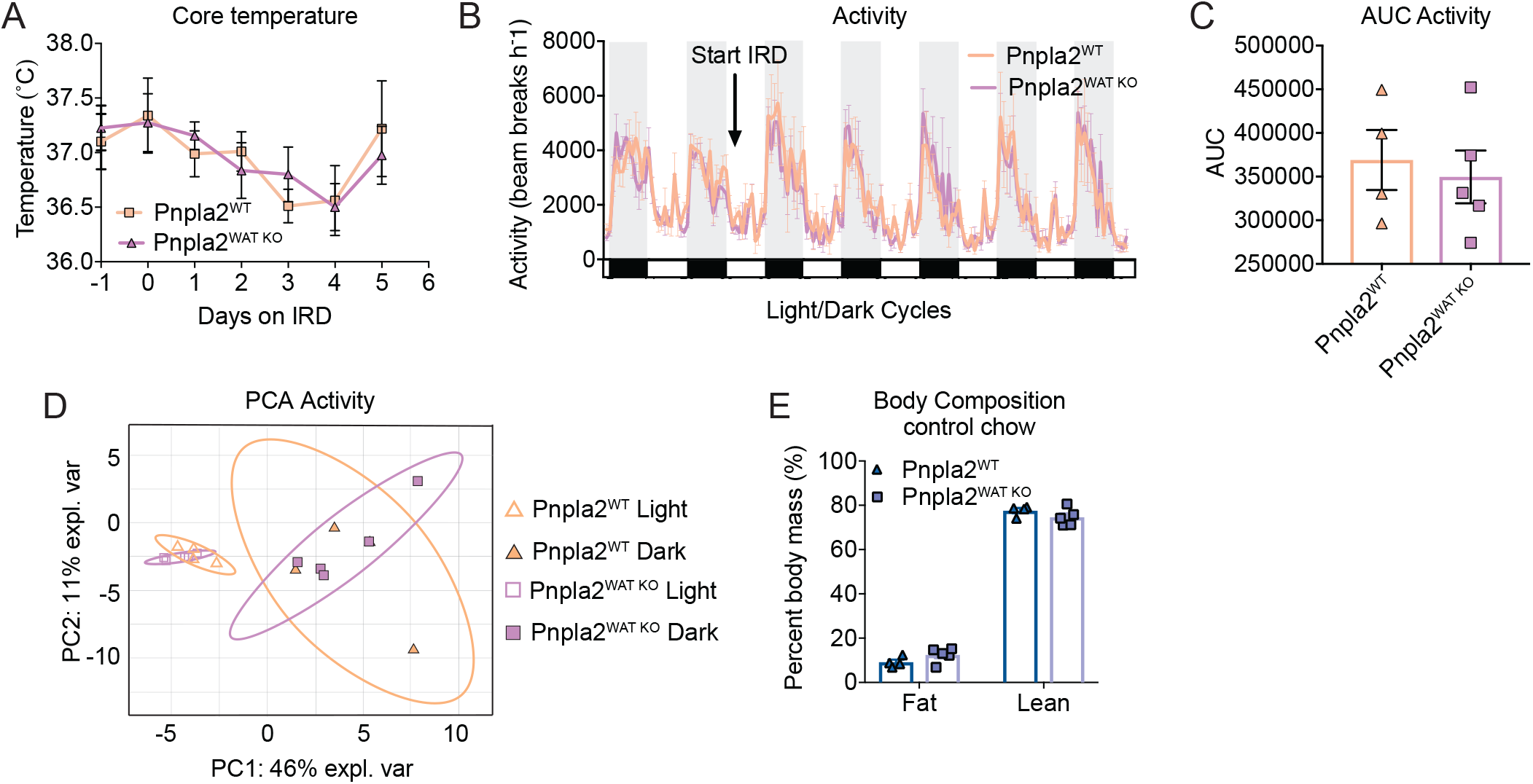
Dietary iron induced adipose-specific ATGL activity protects from wasting of lean energy stores. Littermate *Pnpla2;Fabp4 cre-* (Pnpla2^WT^) and *Pnpla2;Fabp4 cre+* (Pnpla2^WAT KO^) males between six and eight-weeks old were housed in comprehensive laboratory animal monitoring system (CLAMS) metabolic cages for 6 days. Mice were provided 2% carbonyl iron diet (IRD) after brief acclimation period. Daily measurements were taken for body mass, core temperature and food intake. **(A)** Body temperature of Pnpla2^WT^ and Pnpla2^WAT KO^ mice fed control chow prior to IRD supplementation. **(B)** Average hourly activity levels plotted as total X directional beam breaks per hour for Pnpla2^WT^ and Pnpla2^WAT KO^ mice fed IRD and **(C)** Area under the curve analysis for total activity in CLAMS **(D)** Principal component analysis for activity of IRD-fed Pnpla2^WT^ and Pnpla2^WAT KO^ mice in light/dark cycles. Ellipses are indicative of 95% confidence intervals. **(E)** Body composition analyses of Pnpla2^WT^ and Pnpla2^WAT KO^ mice fed control chow prior to IRD supplementation using EchoMRI. Fat and lean mass is normalized to total body mass. All CLAMS data plotted in zeitgeber time. White/black boxes on X axis represent light/dark cycles of 24-hr day. AUC of CLAMS analyses taken from total average values per mouse. Data represent mean ±SEM; Pnpla2^WT^ n=4, Pnpla2^WAT KO^ n=5. Related to **Figure 5**.

